# Rare *SH2B3* coding variants identified in lupus patients impair B cell tolerance and predispose to autoimmunity

**DOI:** 10.1101/2023.04.27.538529

**Authors:** Yaoyuan Zhang, Rhiannon Morris, Ayla May D. Lorenzo, Xiangpeng Meng, Nadia J. Kershaw, Pamudika Kiridena, Grant J. Brown, Gaétan Burgio, Jean Y. Cappello, Qian Shen, Hao Wang, Cynthia M. Turnbull, Tom Lea-Henry, Maurice Stanley, Zhijia Yu, Fiona Ballard, Aaron Chuah, James C. Lee, Ann-Maree Hatch, Alexander P. Headley, Peter Trnka, Dominic Mallon, Jeffery T. Fletcher, Giles D. Walters, Mario Šestan, Marija Jelušić, Matthew C. Cook, Vicki Athanasopoulos, David A. Fulcher, Jeffrey J. Babon, Carola G. Vinuesa, Julia I. Ellyard

## Abstract

Systemic lupus erythematosus (SLE) is a heterogeneous autoimmune disease, with a clear genetic component. While most SLE patients carry rare gene variants in lupus risk genes, little is known about their contribution to disease pathogenesis. Amongst them, *SH2B3* - a negative regulator of cytokine and growth factor receptor signaling – harbors rare coding variants in over 5% of SLE patients. Here we show that unlike the variant found exclusively in healthy controls, most *SH2B3* rare variants found in lupus patients are predominantly hypomorphic alleles. Generation of two mouse lines carrying variants orthologous to those found in patients revealed SH2B3 is important to limit the numbers of immature and transitional B cells. Furthermore, hypomorphic SH2B3 was shown to impair negative selection of immature/transitional self-reactive B cells and accelerate autoimmunity in sensitized mice, at least in part due to increased IL-4R signaling and BAFF-R expression. This work identifies a previously unappreciated role for *SH2B3* in human B cell tolerance and lupus risk.

**Summary:** Zhang *et al*. reveal a role for hypomorphic SH2B3 in lupus risk. The study shows rare and damaging variants identified in lupus patients enable breach of B cell immune tolerance checkpoints and suggests involvement for dysregulated IL-4R signaling and BAFF-R expression.

## Introduction

Systemic lupus erythematosus (SLE) is the prototypic systemic autoimmune disease with diverse clinical manifestations driven by a combination of genetic and environmental factors. Despite its heterogeneous nature, SLE patients share some common features and pathogenic mechanisms. Lupus is characterized by the presence of autoantibodies, especially anti-nuclear antibodies (ANAs) (Tsokos et al., 2016), and deposition of immune complexes (IC) that leads to organ damage (Koffler et al., 1971). Autoantibodies are secreted by autoreactive B cells that evade central and peripheral checkpoints required for establishing self-tolerance (Yurasov et al., 2005). This can be a result of multiple factors including aberrant toll-like receptor (TLR) signaling (Fillatreau et al., 2021), dysregulated cytokines and cytokine receptor signaling (Batten et al., 2000; Granato et al., 2014; Thien et al., 2004), as well as impaired apoptosis and apoptotic cell clearance (Ellyard et al., 2014; Liu et al., 2006; Sisirak et al., 2016).

A role for genetic factors in the pathogenesis of SLE is supported by results from twin concordance studies (Block et al., 1975; Deafen et al., 1992; Ulff-Møller et al., 2018), and >100 susceptibility loci have been identified by genome-wide association studies (GWAS) (Kwon et al., 2019). More recently, whole genome/exome sequencing (WGS/WES) has enabled identification of highly penetrant and damaging rare genetic variants that cause monogenic forms of SLE (Lo, 2016; Omarjee et al., 2019). We previously investigated the presence of rare coding variants in lupus-associated genes – many of them discovered through GWAS - in SLE patients and established that the majority of patients carry one or more such rare variants (Jiang et al., 2019). Functional studies of rare coding variants in *BLK*, *BANK1, P2RY8* and *TLR7* have provided mechanistic insights into disease pathogenesis (Brown et al., 2022; He et al., 2021; Jiang et al., 2019). Our study also revealed that 5.26% of SLE-patients carried rare variants in *SH2B3* (Jiang et al., 2019). Here we describe the impact of these variants on protein function and predisposition to autoimmunity.

*SH2B3* encodes lymphocyte adaptor protein LNK, a negative regulator of numerous cytokine and growth factor receptors transduced by Janus kinases JAK2 (Bersenev et al., 2008) and JAK3 (Cheng et al., 2016) as well as receptor tyrosine kinases c-KIT (Simon et al., 2008) and FLT3 (Lin et al., 2012). Variants in *SH2B3* have been associated with myeloproliferative neoplasms (MPN) (Coltro et al., 2019), idiopathic erythrocytosis (IE) (McMullin et al., 2011) and autoimmune diseases including SLE (Alcina et al., 2010; Bentham et al., 2015; Morris et al., 2016; Wang et al., 2021), rheumatoid arthritis (RA) (Okada et al., 2014), type 1 diabetes (T1D) (Steck et al., 2017), and multiple sclerosis (MS) (Alcina et al., 2010). A 2013 study on two siblings with homozygous *SH2B3* variant D231Profs*38 reported the development of precursor B-cell acute lymphoblastic leukemia (ALL) with Hashimoto thyroiditis and suspected autoimmune hepatitis (Perez-Garcia et al., 2013), while a recent case report described a novel clinical syndrome involving myelopreliferation and multi-organ autoimmunity in unrelated patients carrying homozygous *SH2B3* variants R148Profs*40 and V402M (Blombery et al., 2022). *Sh2b3^-/-^* mice display dysregulation of numerous hematopoietic cell types including hematopoietic stem cells (HSCs) (Bersenev et al., 2008), B and T lymphocytes (Katayama et al., 2014; Mori et al., 2014; Takaki et al., 2000), platelets (Takizawa et al., 2010), dendritic cells (DCs) (Mori et al., 2014) and neutrophils (Laroumanie et al., 2018). Although *SH2B3* variants have been associated with autoimmune diseases, the cellular mechanisms by which they contribute to autoimmune pathogenesis are yet to be elucidated.

Using *in vitro* assays and mouse models engineered with CRISPR/Cas9 to harbor patient *Sh2b3* variants, we present data showing that rare *SH2B3* variants act as hypomorphic alleles that impair B cell tolerance mechanisms predisposing to autoimmunity.

## Results

### Loss-of-function *SH2B3* variants in SLE patients

Previous bioinformatic analysis of WGS/WES in our cohort of 132 SLE patients and 97 healthy controls revealed that 5.26% and 2.06% of patients and healthy controls respectively carried rare (minor allele frequency, MAF < 0.005) *SH2B3* coding variants. Clinical information for the patients is shown in **Supplementary Table 1**. Additional analyses for rare variants in 22 genes that can cause human SLE when mutated (**Supplementary Table 2**) revealed that one patient (D.II.1) was compound heterozygous for *ACP5* (a pathogenic variant p.G109R (Lausch et al., 2011) and a novel variant p.C238R). This patient also had a variant of unknown significance in *IFIH1* p.A542E. Another patient (A.II.1) was heterozygous for a variant of unknown significance in *ADAR*, p.I939V. No additional mutations in known SLE-causing genes were identified in the other patients. Of the seven rare heterozygous *SH2B3* variants identified, five (C133Y, E400K, A536T, Q540X, R566Q) were exclusive to SLE kindreds and one (E208Q) was found in both SLE and HC. The seventh variant (R43C) was identified only in a heathy control (**Figure 1A**). Whereas the SLE-associated variants occurred in the PH domain (E208Q), SH2 domain (E400K) and near the C-terminus of the protein (**Figure 1B**), the R43C variant identified in the healthy control localized to the dimerization domain.

**Figure 1.**
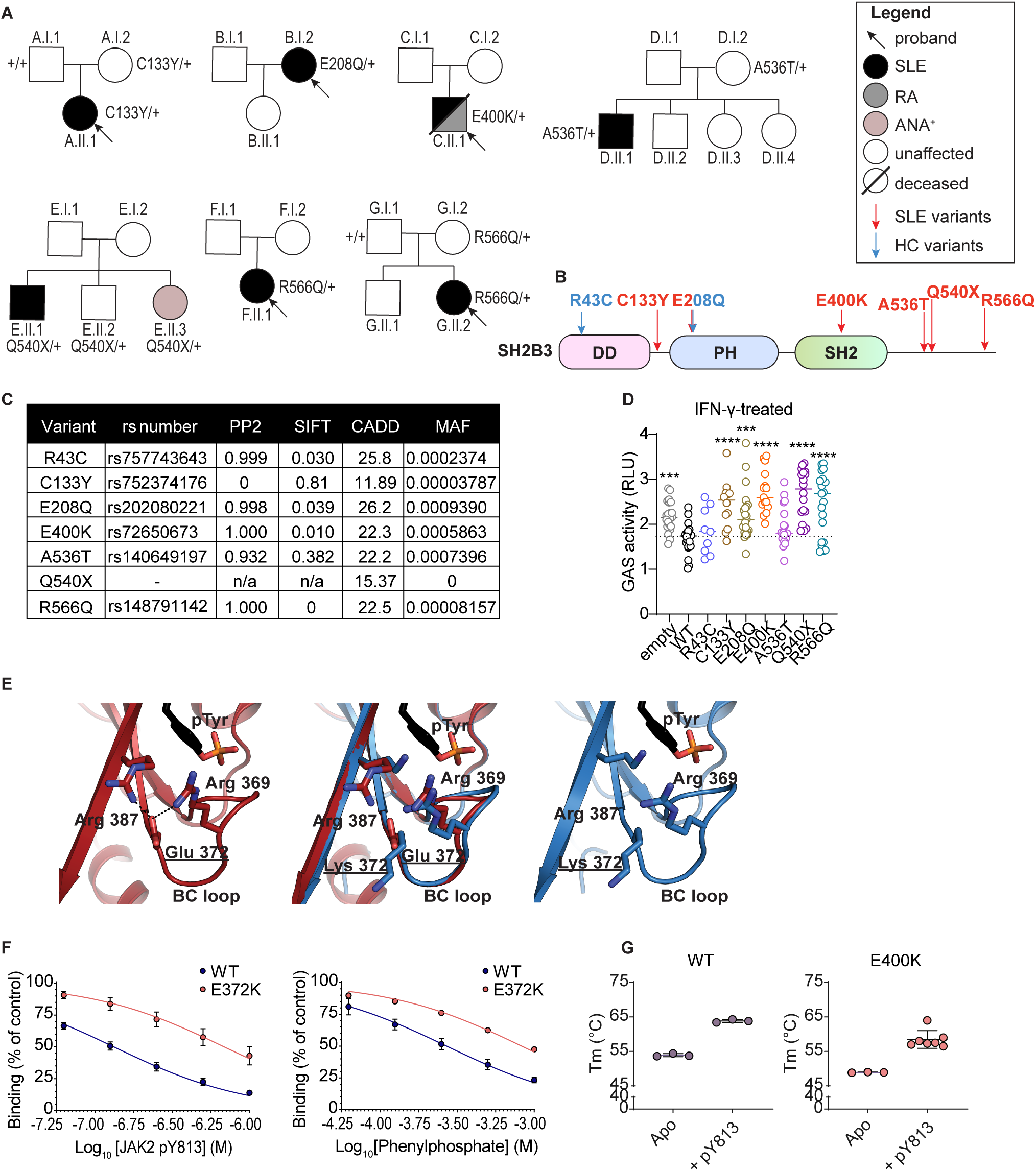
Rare *SH2B3* variants identified in SLE patients and healthy controls (HCs). (**A**) Pedigrees of families with rare single nucleotide variants (SNV) in *SH2B3* associated with autoimmunity. Black arrows indicate the proband of each cohort. Individuals colored black are affected by SLE, individuals colored grey are affected by rheumatoid arthritis (RA). Individuals with question marks are not yet sequenced. The amino acid changes in SH2B3 protein are shown next to sequenced individuals, with “+” sign indicating “wildtype”. (**B**) Location of patient specific and healthy control (HC) SNVs in the schematic diagram of SH2B3 amino acid sequence. DD: dimerization domain, SH2: Src homology 2 domain, PH2: Pleckstrin homology domain. (**C**) Summary of variant information, *in silico* deleteriousness prediction and minor allele frequency of each SNV from healthy control and SLE patients. PP2: PolyPhen-2. (**D**) Relative activity of STAT1-GAS in HEK293 cells overexpressing empty vector, wildtype or mutant human *SH2B3* in presence of IFN-γ (50 ng/mL) stimulation. Relative activity was calculated by normalizing all STAT1-GAS-Firefly/CMV-Renilla ratios to the average ratios in the unstimulated cells transfected with empty vectors. Results were pooled from seven independent experiments. Means are shown as bars and all conditions were compared to cells transfected with WT SH2B3. Linear mixed-effect model (lmer) with estimated marginal means (emmeans) using experiment as the blocking factor was used for the statistical analyses. Significance levels are indicated with asterisks. *: p < 0.05, **: p < 0.01, ***: p < 0.001, ****: p < 0.0001. (**E**) Hydrogen bonding network between Arg369-Glu372-Arg387 (left) that helps position the BC loop relative to the central β-sheet and disrupted hydrogen bond formation (right) when Glu372 is replaced with Lys. (**F**) Percentage of binding of murine WT or E372K SH2B3 to IL6ST pY757 peptide chip after pre-incubation with varying concentrations of mutant JAK pY813 peptides (left) or phenylphosphate (right). Data are shown as the mean ± SD of technical replicates from at least three independent experiments. (**G**) Melting temperatures of WT (left) and E400K (right) SH2B3 protein at apo state (unbound) or bound to pY813 JAK2. Data are displayed as the mean ± SD of technical replicates from at least three independent experiments.

We asked whether the rare variants observed in SLE patients conferred functional defects. All variants, except the novel nonsense mutation Q540X, were predicted to be damaging by at least one *in silico* prediction algorithm (**Figure 1C**). To assess protein function, we examined IFN-γR signaling that is in part transduced by JAK2, a target of SH2B3 negative regulation (Igarashi et al., 1994). While overexpression of WT SH2B3 was able to partially suppress STAT1-mediated GAS activity in HEK293 cells stimulated with IFN-γ, this regulation was lost in cells overexpressing all rare SLE patient variants, except A536T (**Figure 1D, Figure S1A**). The A536T variant was present in a proband that also carried biallelic variants in *ACP5*, which is a likely cause of the patient’s disease (Kara et al., 2020). GAS activity was also suppressed in cells expressing the SH2B3 R43C variant present in the healthy control. Of the SLE patient variants, Q540X, R566Q and E400K exerted the strongest functional defects.

We were particularly interested in the variant E400K, as it is a key residue within the SH2 binding pocket that binds phosphotyrosine (pTyr) residues of target proteins (Morris et al., 2021). E400 forms part of a hydrogen bonding network that appears to be required for the stability of the BC loop that forms the base of the pTyr binding pocket. To confirm the impaired function observed in the luciferase assays, we performed structural analysis of the murine SH2B3 SH2 domain containing the orthologous E372K mutation (**Supplementary Table 3)**. The murine and human SH2B3 SH2 domains share 92.9% identity with high conservation across peptide binding residues. Co-crystallization with the JAK2 pY813 motif showed that, whilst the motif could still bind, substitution of the glutamate with a lysine resulted in the loss of the hydrogen bonding network with Arg 369 and Arg 387, potentially destabilizing the BC loop (**Figure 1E;** WT: **PDB ID 7R8W,** E372K: **PDB ID: 8CZ9**). Supporting this hypothesis, a decrease in the affinity of the SH2B3/pY813 interaction was observed using a competition assay (E372K IC_50_ = 685 n*M* compared to WT = 127 n*M*), as was the affinity for the pTyr mimetic, phenyl phosphate (**Figure 1F**). The reduction in affinity for JAK2 pY813 was due to a slower on-rate, rather than a compromised off-rate (**Figure S1B**).

In addition, the E372K SH2 domain showed decreased thermal stability compared to WT SH2 domain both alone (apo form) or when bound to JAK2 pY813 (**Figure 1G**) or other peptides (**Figure S1C**). These findings suggest that E400K SH2B3 impairs protein function as a result of both reduced affinity to pTyr targets and reduced stability due to the loss of the hydrogen bonding network that requires Glu372. Together, these results indicate that the majority of *SH2B3* variants found in SLE patients are hypomorphic alleles.

### SLE patient *SH2B3* variants behave as hypomorphs in mice

To understand whether these variants conferred phenotypes *in vivo*, we used CRISPR/Cas9 gene editing technology to generate two mouse models, one carrying the E400K orthologue E372K allele (*Sh2b3^E372K^*) that was shown to be important in pTyr binding of the SH2 domain, and another carrying the R566Q orthologue R530Q allele (*Sh2b3^R530Q^*) that showed profound loss of activity (**Figure 1D**) and was identified in two unrelated patients (**Figure 1A**). A third strain containing a 2-base-pair deletion (*c.1107_1108del,* coding RefSeq NM_001306127 or *p.G371RfsTer56*) causing a frameshift and premature stop codon expected to result in a null protein was also generated for comparison.

Previous studies have indicated that Sh2b3-deficient mice exhibit defects in hematopoiesis especially lymphopoiesis (Velazquez et al., 2002). Consistent with those reports, we observed a large increase in peripheral blood leukocyte and lymphocyte counts in Sh2b3-deficient mice (*Sh2b3^Δ^*), with a similar but weaker increase in *Sh2b3^E372K^* and *Sh2b3^R530Q^*strains, which was statistically significant only in homozygous mice (**Figure 2A, B**). Homozygous mice from all three strains also presented with mild splenomegaly (**Figure 2C**) and increased spleen cellularity (**Figure 2D**). Next, we examined splenic B and T cells and observed that although frequencies were unchanged (**Figure S1D-F**) the total number of both B and T cells increased (**Figure 2E, G**) consistent with the overall increase in lymphocyte counts in the spleen and blood. Furthermore, mixed BM chimeras indicated that *Sh2b3^E372K/E372K^*B and T cells displayed a competitive advantage over *Sh2b3^+/+^*cells in repopulating sub-lethally irradiated *Rag1^-/-^* mice (**Figure 2F, H**).

**Figure 2.**
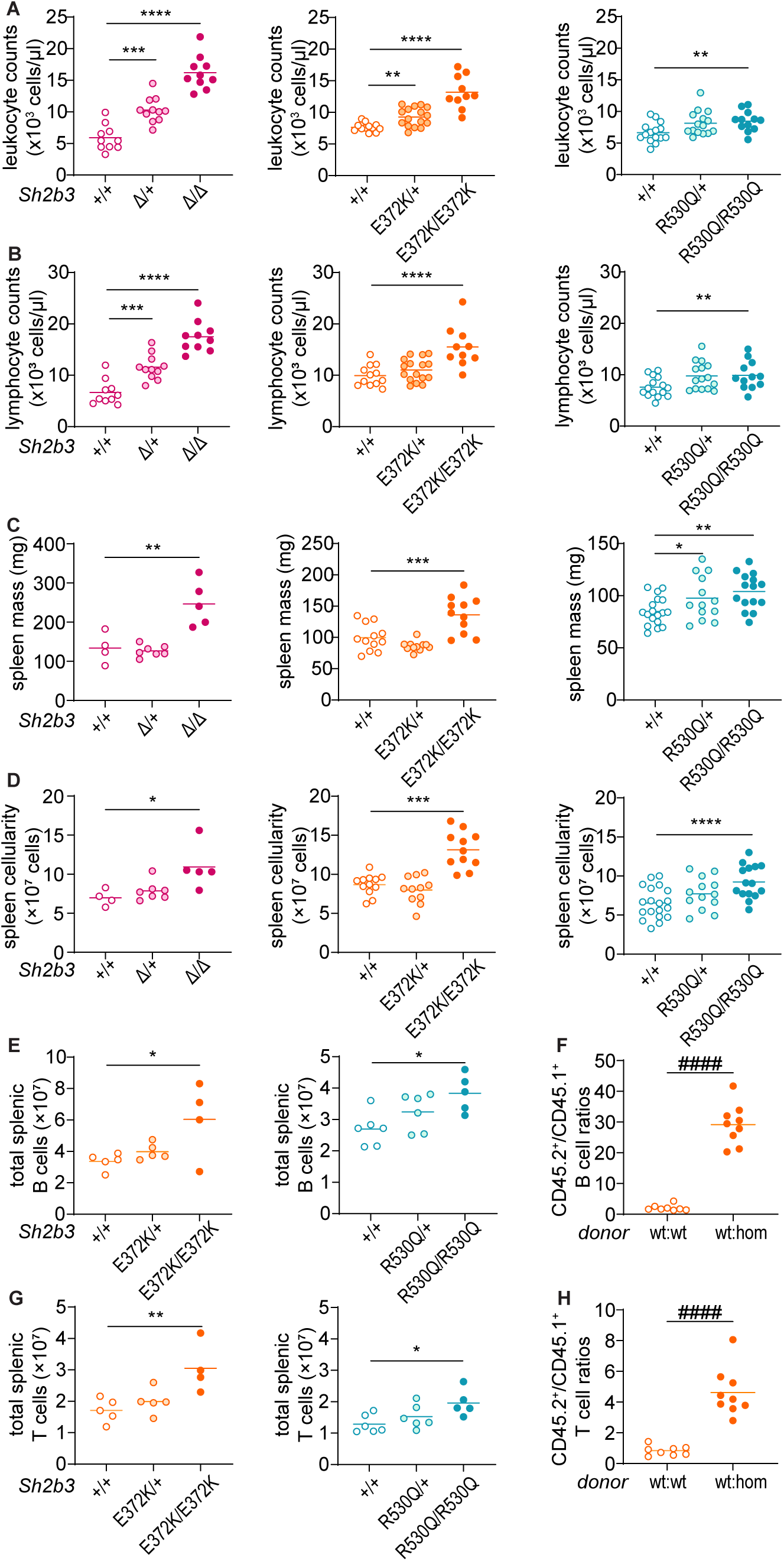
Gross phenotypes of *Sh2b3^Δ^, Sh2b3^E372K^* and *Sh2b3^R530Q^*mice. Peripheral blood (**A**) leukocyte and (**B**) lymphocyte counts determined by ADVIA hematological analyzer. (**C**) Spleen mass and(**D**) cellularity. Total splenic (E) B cells in *Sh2b3^Δ^*, *Sh2b3^E372K^*and *Sh2b3^R530Q^* mice. (**F**) Ratios of CD45.2^+^/CD45.1^+^ B cell percentages in the spleens of 50:50 BM chimeras from CD45.1-*Sh2b3^+/+^*and CD45.2-*Sh2b3^+/+^*/*Sh2b3^E372K/E372K^* mice. (**G**) Total splenic T cells in *Sh2b3^Δ^*, *Sh2b3^E372K^*and *Sh2b3^R530Q^* mice. (**H**) Ratios of CD45.2^+^/CD45.1^+^ T cell percentages in the spleens of 50:50 BM chimeras from CD45.1-*Sh2b3^+/+^* and CD45.2-*Sh2b3^+/+^*/*Sh2b3^E372K/E372K^* mice. Results in A-D were pooled from 2-4 independent experiments. Means are shown as bars. Results in E and G are representative of two independent experiments. Means are shown as bars. Results in Fand H are representative of two independent experiments. Linear mixed-effect (lmer) model ANOVA with multiple comparison and Tukey’s correction and was used for statistical analysis in A-D. Statistical significance is indicated by asterisks. One-way ANOVA with multiple comparison and Dunnett’s correction was used for statistical analysis in E and G. Significance levels are indicated with asterisks. Student-t test was used for statistical analysis in F and H. Significance levels are indicated with hashes. Significance level criteria described as follow, */#: p < 0.05, **/##: p < 0.01, ***/###: p < 0.001, ****/####: p < 0.0001.

### *Sh2b3^E372K^* and *Sh2b3^R530Q^* mice show dysregulated B cell development

Previous studies on *Sh2b3^-/-^* mice have reported accumulation of transitional B cells in the spleen, as well as pro- and pre-B cells in the BM due to unrestrained c-KIT and IL-7R signaling (Cheng et al., 2016; Takaki et al., 2000). As expected, we observed increased frequencies of splenic transitional B cells in *Sh2b3^Δ/+^* and *Sh2b3^Δ/Δ^* mice (**Figure 3A, B**). We also observed increased frequency and total numbers of transitional B cells in both *Sh2b3^E372K/E372K^* and *Sh2b3^R530Q/R530Q^* mice (**Figure 3A, B, Figure S1G**). This increase was due to B cell-intrinsic Sh2b3 signalling (**Figure 3C**), and occurred primarily within the transitional 1 (T1) B cell compartment (**Figure 3D, E, Figure S1G-I**) with a milder increase in transitional 2 (T2) and transitional 3 (T3) B cells (**Figure S1J-P**).

**Figure 3.**
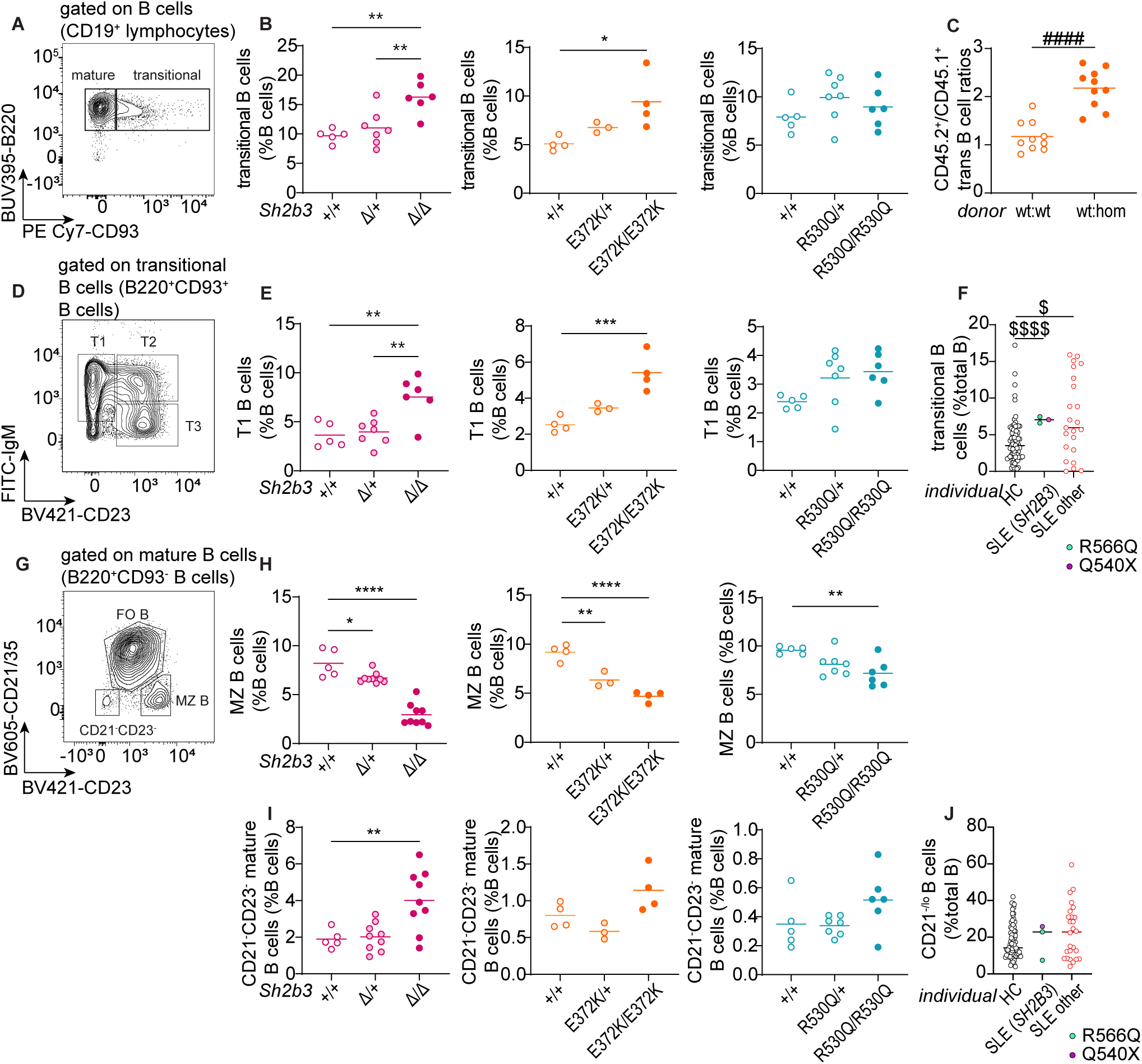
B cell phenotypes in the spleens of *Sh2b3^Δ^*, *Sh2b3^E372K^* and *Sh2b3^R530Q^*mice, and peripheral blood of SLE patients carrying *SH2B3* variants R566Q and Q540X. (**A**) Representative flow cytometric plot showing the gating of mature (B220^+^CD93^-^) and transitional (B220^+^CD93^+^) B cells. (**B**) Dot plots showing the percentages of splenic transitional B cells in *Sh2b3^Δ^* (left), *Sh2b3^E372K^* (middle) and *Sh2b3^R530Q^* (right) mice. (**C**) Ratios of CD45.2^+^/CD45.1^+^ transitional B cell percentages in the spleens of 50:50 BM chimeras from CD45.1-*Sh2b3^+/+^* and CD45.2-*Sh2b3^+/+^*/*Sh2b3^E372K/E372K^* mice. (**D**) Representative flow cytometric plot showing the gating of T1 (IgM^+^CD23^-^), T2 (IgM^+^CD23^+^) and T3 (IgM^-^CD23^+^) B cells. (**E**) Dot plots showing the percentages of splenic T1 B cells in *Sh2b3^Δ^* (left), *Sh2b3^E372K^*(middle) and *Sh2b3^R530Q^* (right) mice. (**F**) Percentages of human transitional B cells in the peripheral blood of healthy controls (HC, n = 77), SLE patients carrying *SH2B3* variants (R566Q in turquoise, n = 2, and Q540X in purple, n = 1) and other SLE patients (n = 23). (**G**) Representative flow cytometric plot showing the gating of CD21^-^CD23^-^ mature B cells and marginal zone (MZ) B cells (CD21/35^-^CD23^+^). (H) Dot plots showing the frequencies of MZ B cells as percentages of total splenic B cells in *Sh2b3^Δ^* (left), *Sh2b3^E372K^* (middle) and *Sh2b3^R530Q^* (right) mice. (I) Frequencies of CD21^-^CD23^-^ mature B cells as percentages of total splenic B cells in *Sh2b3^Δ^* (left), *Sh2b3^E372K^* (middle) and *Sh2b3^R530Q^*(right) mice. (**J**) Percentages of human CD21^lo/-^ B cells in the peripheral blood of healthy controls (HC, n = 77), SLE patients carrying *SH2B3* variants (R566Q in turquoise, n = 2, and Q540X in purple, n = 1) and other SLE patients (n = 23). Results are representative of three independent experiments while means are shown as bars in B, E, G and I. One-way ANOVA with multiple comparison and Dunnett’s correction was used for statistical analysis in B, E, H and I. Significance levels are indicated with asterisks. Student-t test was used for statistical analysis of chimera data in C. Significance levels are indicated with hashes. Brown-Forsythe and Welch ANOVA tests were used for statistical analysis in F and J. Dollar signs are used to indicate significance levels. Significance level criteria described as follow, */#/$: p < 0.05, **/##/$$: p < 0.01, ***/###/$$$: p < 0.001, ****/####/$$$$: p < 0.0001.

Whilst the frequency of mature B cells was decreased in *Sh2b3^E372K/E372K^*and *Sh2b3^R530Q/R530Q^* mice, total numbers remained elevated as a result of the overall increase in splenic B cells (**Figure S2A-D**). Marginal zone (MZ) B cells in both *Sh2b3^E372K/E372K^*, *Sh2b3^R530Q/R530Q^* and *Sh2b3^Δ/Δ^* mice were also decreased, with heterozygous *Sh2b3^E372K/+^* and *Sh2b3^R530Q/+^*displaying an intermediate phenotype (**Figure 3G, H**). This change was not due to a reduction of MZ B cell numbers (**Figure S2E**), but an increase in the other mature B cell subsets (**Figure S2D, F**). Analysis of 50:50 mixed BM chimeras revealed that the decrease in MZ B cells was cell-intrinsic (**Figure S2H**). This phenotype has not previously been reported in *Sh2b3^−/−^* mice but loss of MZ B cells; although a decreased in MZ B cells has been observed in lupus-prone B6.*Yaa* mice, this was cell-extrinsic as result of enhanced activation by antigen-bearing DCs (Santiago-Raber et al., 2010). We also examined CD21/35^-^CD23^-^ mature B cells, a B cell subset enriched in atypical memory B cells (ABCs), that are found to be elevated in scenarios of chronic antigen stimulation such as patients with autoimmunity (Rubtsov et al., 2011; Wehr et al., 2004). There was a slight but not statistically-significant cell-extrinsic increase of CD21/35^-^CD23^-^ mature B cells in both *Sh2b3^E372K/E372K^*, *Sh2b3^R530Q/R530Q^* and *Sh2b3^Δ/Δ^* mice (**Figure 3I, Figure S2J**). No difference in ABCs was observed (**Figure S2K-M**).

Phenotyping of the three human SLE patients with rare variants in *SH2B3* (one carrying Q540X and two carrying R566Q), (**Figure 1A)** using our established protocols (Ellyard et al., 2019) revealed a higher frequency of transitional B cells compared to healthy controls, which is also observed in other SLE patients (**Figure 3F**) and reported previously (Sims et al., 2005). CD21^-/lo^ B cells, a population also reported to be elevated in SLE patients (Wehr et al., 2004), showed a trend of increase in two SLE patients carrying rare *SH2B3* variants and other SLE patients (**Figure 3J**).

Having established a dysregulation in peripheral B cells, we investigated B cell development in the BM. Consistent with reports from *Sh2b3*^-/-^ mice (Takaki et al., 2000), we observed a significant increase of B cell precursors as a percentage of BM B cells in *Sh2b3^E372K/E372K^*mice and a similar trend in *Sh2b3^R530Q/R530Q^* mice (p=0.0536) (**Figure S3A, B**). Among B cell precursors, there was a cell-intrinsic increase of pre-B cell percentages in both strains (**Figure S3C-E**), but no consistent differences in pre-pro B or pro-B cells (**Figure S3F, G**). We also identified a trend of increased immature B cells (**Figure S3H**) and a significant decrease in mature B cells (**Figure S3I**). Our results are consistent with the published data on *Sh2b3^-/-^* mice and suggest that impaired negative regulation of growth factor/cytokine receptor signaling in BM B cells causes increased B cell generation and therefore accumulation in the BM.

### Increased T cell activation in *Sh2b3^E372K^* and *Sh2b3^R530Q^* mice

Studies in human cell lines have suggested that SH2B3 may play important roles in T cell activation by negatively regulating components of the TCR signaling pathway through interactions with PLCγ, GRB2, PI3K and FLNA (He et al., 2000; Huang et al., 1995; Takaki et al., 1997), and indirectly repress T cell activation through cytokine receptor signaling including IL-7R and IL-15R in dendritic cells (Katayama et al., 2014; Lawson et al., 2015). We observed decreased CD4/CD8 T cell ratios in the peripheral blood of *Sh2b3^E372K/+^, Sh2b3^E372K/E372K^* (**Figure S 4A, B**) and spleens of *Sh2b3^R530Q/R530Q^* mice (**Figure S 4C**), but not in the peripheral blood of *Sh2b3^R530Q^*mice (**Figure S 4B**) and only a slight trend in the spleens of *Sh2b3^E372K/E372K^*mice (**Figure S 4C**). Such phenotype has been reported among SLE patients (Maeda et al., 1999). *Sh2b3^E372K/+^* and *Sh2b3^E372K/E372K^*, but not *Sh2b3^R530Q^* mice, also presented with increased percentages of double negative (DN: CD4^-^CD8^-^) T cells in peripheral blood (**Figure S 4D**). This subset has also been found to be expanded in some SLE patients (Crispín et al., 2008), although it was not present in the SLE patients with *SH2B3* rare variants (data not shown). Within CD4 T cells, we observed increased percentages and total numbers of effector memory CD4 T (CD4 T_EM_) cells and regulatory T (T_reg_) cells in the spleens of *Sh2b3^E372K/+^* and *Sh2b3^R530Q/R530Q^*mice (**Figure S 4E-I**). T_reg_ suppressive function was normal (**Figure S 4J**). Among CD8 T cells, we observed increased percentages of CD8 T effector memory (T_EM_) cells in the peripheral blood of *Sh2b3^E372K/+^* and *Sh2b3^E372K/E372K^*mice (**Figure S 4K, L**) and the spleens of *Sh2b3^R530Q/R530Q^*mice (**Figure S 4M, N**). Analysis of 50:50 mixed BM chimeras revealed that these T cell phenotypes were largely cell-extrinsic (**Figure S 4O-T**). Together, our findings reveal a trend towards increased activated and/or effector T cells in mice carrying hypomorphic *Sh2b3* variants.

### Sensitized *Sh2b3^E372K^* mice manifest accelerated autoimmunity

Having demonstrated that *SH2B3* variants identified in SLE patients are hypomorphic alleles resulting in immune dysregulation, we next asked whether they confer susceptibility to autoimmunity. A hallmark of SLE is production of anti-nuclear antibodies (ANAs), particularly against dsDNA. We analyzed levels of anti-nuclear antibodies (ANAs) and anti-DNA antibodies in the sera of *Sh2b3^E372K^* mice but found no substantial differences compared to wildtype littermate controls (**Figure A, B**). Hence, our results indicate that *Sh2b3^E372K^* mice do not develop autoimmunity spontaneously.

We then considered whether autoimmunity may be accelerated or more severe in a sensitized background. Pristane, a hydrocarbon compound derived from shark liver oil and mineral oil (Ackman, 1971), also known as 2,6,10,14-tetramethylpentadecane (TMPD), is a well-established inducer of lupus-like disease when administrated intraperitoneally (i.p.) in mice (Reeves et al., 2009). We injected 12-week-old *Sh2b3^E372K^*mice i.p. with either pristane or PBS control and monitored development of anti-DNA autoantibodies in the serum (**Figure C**). *Sh2b3^E372K/E372K^*mice developed significantly higher titers of anti-DNA IgG 10 weeks post pristane treatment (**Figure D, E**). However, this difference was lost by 20 weeks when pristane-treated *Sh2b3^+/+^, Sh2b3^E372K/+^*and *Sh2b3^E372K/E372K^* mice all displayed similar titers of anti-DNA autoantibodies (**Figure F**). Pristane-treatment also caused IgG IC deposition in the glomeruli of mice at 24 weeks (**Figure S5A, B**), with some mice also displaying minimal glomerular leukocyte infiltration (score 1 out of 4) (**Figure S5D**). We did not observe an increase in proteinuria in pristane-treated mice compared to the PBS treatment group (data not shown). Anti-DNA IgG titers did not change significantly over time in PBS-treated control mice regardless of genotype (**Figure G-I**) with few mice displaying glomerular IgG IC deposition (**Figure S5A, C**) or minimal glomerular infiltration (**Figure S5E**).

### Decreased SH2B3 function compromises B cell tolerance at multiple immune checkpoints

The fact that sensitizing *Sh2b3^E372K/E372K^* mice led to accelerated development of autoantibodies together with the increase in immature/transitional B cells that are known to harbor autoreactivity (Wardemann et al., 2003), suggest that hypomorphic *Sh2b3* variants promote the escape of autoreactive B cells. To examine this, we took advantage of the SW_HEL_-mHEL^3×^ mixed chimera model, in which bone marrow (BM) with B cells expressing transgenic immunoglobulin heavy and light chains specific for hen egg lysozyme (HEL) from SW_HEL_ mice, is engrafted into irradiated mHEL^3×^ mice that express HEL as a membrane-bound self-antigen (Burnett et al., 2018). We crossed the *Sh2b3^E372K^* mice to SW_HEL_ mice (hereinafter referred to as SW_HEL_-*Sh2b3^E372K^* mice) and transplanted BM cells from these crosses into irradiated WT (non-self-reactive) or mHEL^3×^ mice (self-reactive) recipients (**Figure A**). Consistent with previous reports (Lau et al., 2019), we observed an 80-90% decrease in the frequency of HEL-specific mature *Sh2b3^+/+^* (wild-type) B cells in the BM of mHEL^3x^ recipients compared to wild-type recipients; evidence that autoreactive HEL-binding B cells are purged from the recirculating B cell repertoire (**Figure B, C**). In contrast, no decrease in self-reactive B cells was observed in recipients of HEL-binding *Sh2b3^E372K/E372K^*BM, suggesting that reduced SH2B3 function prevents removal of autoreactive B cells. Interestingly, there appeared to be partial negative selection of HEL-binding *Sh2b3^E372K/+^* B cells, indicating an intermediate phenotype. No difference was seen in non-HEL binding mature B cells, indicating that purging of WT B cells observed in HEL^3x^ recipients is antigen-specific (**Figure S5F**).

To determine at which immune checkpoint tolerance is initially breached, we first looked at B cells developing in the BM. Autoreactive B cell clones are typically removed through negative selection at the pre-B (the first stage in B cell development at which pre-BCR are expressed on most B cells) to immature development stage. Similar to mature B cells, there was a failure to delete immature *Sh2b3^E372K/E372K^* HEL-specific B cells in HEL^3x^ recipients, although this was not observed for B cells heterozygous for the *Sh2b3^E372K^* allele (**Figure D**). In contrast, HEL-specific pre-B cells were not deleted in mHEL^3x^ recipients, consistent with these cells not having undergone negative selection (**Figure E**), and in fact we observed an increase in the mHEL^3x^ autoreactive environment regardless of *Sh2b3* genotype.

We next examined immune tolerance checkpoints in the spleen, where autoreactive transitional B cells are negatively selected and inhibited from entering the mature follicular B cell pool. We observed a failure to remove HEL-specific transitional *Sh2b3^E372K/E372K^*B cells in mHEL^3×^ recipients (**Figure F**), with heterozygous *Sh2b3^E372K/+^* B cells leading to an intermediate phenotype. This same enrichment in self-reactive B cells was also observed amongst HEL-specific MZ and follicular (FO) B cells (**Figure G-I**) in mHEL^3×^ mice receiving SW_HEL_-*Sh2b3^E372K/E372K^ or Sh2b3^E372K/+^*BM, whereas these compartments were significantly devoid of self-reactive B cells in mHEL^3×^ mice receiving SW_HEL_-*Sh2b3^+/+^* BM. This indicates a specific defect in the tolerance checkpoint between splenic transitional and mature B cells. Downregulation of IgM surface expression is an indicator of recent BCR engagement and thus often used as a measure of B cell anergy (Goodnow et al., 1989) although it can also indicate activation. Consistently, IgM expression in the chimeras was decreased on HEL-specific *Sh2b3^+/+^* transitional and follicular (FO) B cells in mHEL^3×^ recipient mice compared to WT recipients (**Figure S5K, L**). Similarly, IgM was significantly downregulated in HEL-specific S*h2b3^E372K/E372K^*FO B cells in the mHEL^3×^ recipient environment, IgM downregulation on transitional S*h2b3^E372K/E372K^* B cells was less affected. Interestingly, as previously noted (Lau et al., 2019), an autoreactive environment does not appear to modulate IgM expression in MZ B cells regardless of genotype (**Figure S5M**).

Given this escape of autoreactive S*h2b3^E372K/E372K^* B cells into the mature recirculating pool, we next examined whether they were also activated in the periphery to become an autoantibody-secreting cell; an additional checkpoint in B cell tolerance. Initially we examined the generation of HEL-specific antibodies. We did not observe an increase in serum HEL-specific IgM antibodies in the autoreactive mHEL3x background (**Figure S5N**), whilst S*h2b3^E372K/E372K^* B cells produced less HEL-specific IgG antibodies in mHEL 3x mice compared to wildtype recipients (**Figure S5O**). Interestingly, we did observe a propensity for *Sh2b3^E372K/E372K^* B cells to differentiate ABCs that can be activated to produce pathogenic autoantibodies (Rubtsov et al., 2011; Wang et al., 2018). Whilst, *Sh2b3^+/+^* and *Sh2b3^E372K/+^* B cells appeared to be prevented from differentiating into ABCs in the mHEL^3x^ recipient environment, *Sh2b3^E372K/E372K^*B cells gave rise to a higher frequency of HEL-specific ABCs, suggesting they may not be as effectively removed (**Figure J**). A similar trend was observed for the parent population CD21^-^CD23^-^ mature B cells (**Figure S5Q**). However, the amount of HEL-specific ABC or CD21-CD23-mature B cells was generally low, compared to non-HEL-specific populations (**Figure S5P, R**).

To determine whether changes in receptor editing in SH2B3 hypomorphic B cells may contribute to the breach of central tolerance, we stained for immunoglobulin κ and λ light chains in BM B cells from *Sh2b3^+/+^*, *Sh2b3^E372K/+^* and *Sh2b3^E372K/E372K^* mice but did not observe any difference in the Igκ/Igλ ratios between genotypes for either immature or mature B cells (**Figure 5K, L**). Changes in BCR signalling strength may also influence B cell negative selection (Lam et al., 1997; Nemazee and Bürki, 1989). We performed a calcium flux assay to determine if the E372K allele could regulate BCR signal strength, but observed no difference in the level of calcium flux for *Sh2b3^E372K/+^*or *Sh2b3^E372K/E372K^* transitional or mature B cells compared to *Sh2b3^+/+^* controls (**Figure 5M, N**).

**Figure 4.**
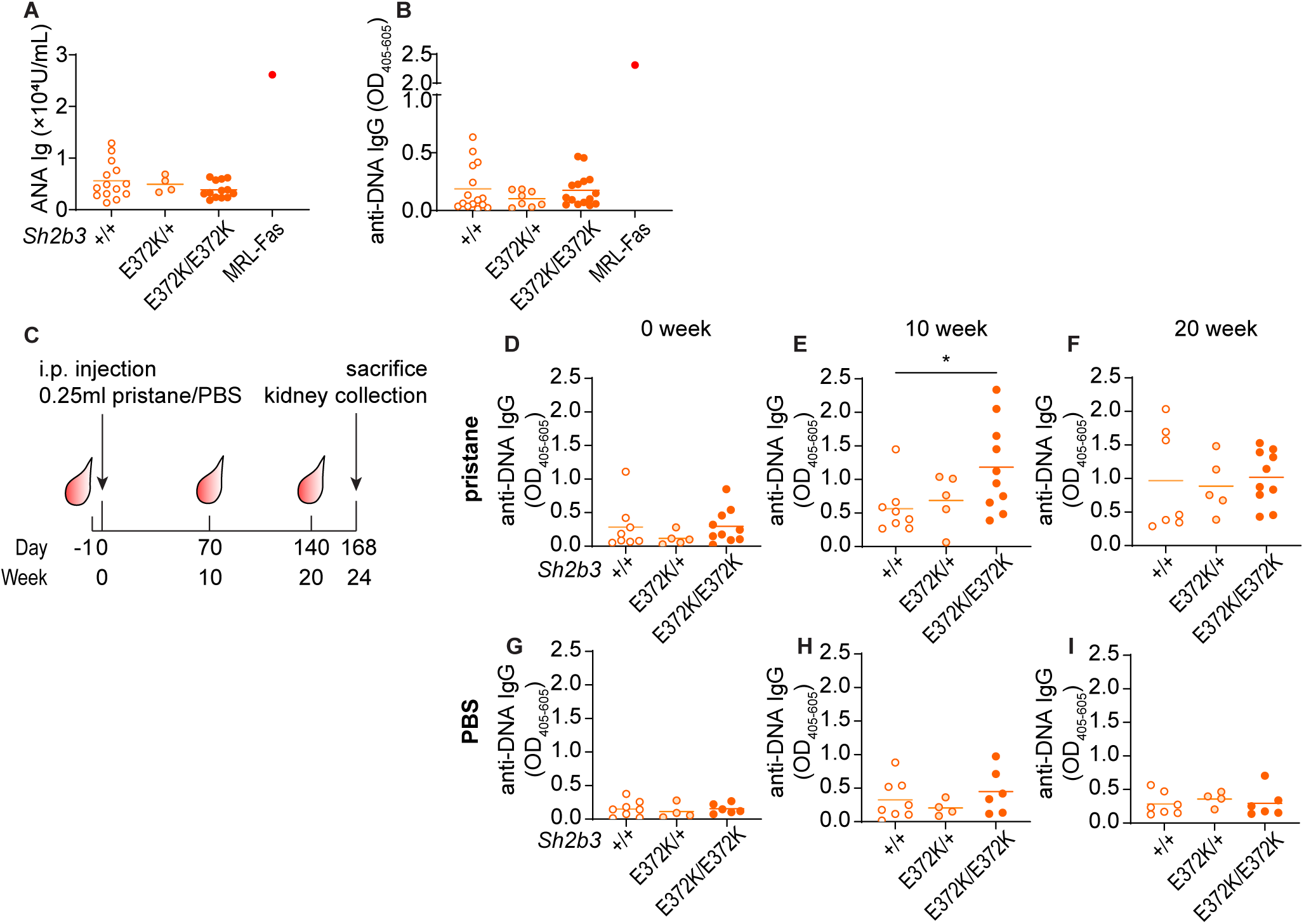
Autoimmune triggers induced exacerbated autoimmunity in *Sh2b3^E372K^* mice. **A-B**. Serum anti-nuclear antibodies (ANAs; **A**) and anti-DNA IgG (**B**) determined by ELISA. Data in a and b are from single experiments. (**C**) Experimental plan of pristane-induced lupus-like autoimmune model in *Sh2b3^E372K^* mice. 14-week-old female *Sh2b3^E372K^* mice were bled one day before the intraperitoneal (i.p.) injection of 0.25-mL pristane or PBS (control) and later 2-, 10- and 20-week post-injection and sacrificed 24-week post-injection for spleen and kidney analysis. Levels of serum anti-DNA IgG in pristane-treated mice at 0 (**D**), 10 (**E**) and 20 (**F**) weeks post treatment, and PBS-treated mice at 0 (**G**), 10 (**H**) and 20 (**I**) weeks post treatment. Means are shown as bars in all dot plots. Data in D-I were pooled from two independent experiments. Fractions of IgG deposit positive and negative kidney samples collected 24-wells after (F) pristane and (G) PBS *i.p.* injections and (H) corresponding representative microscopy images. Green channel: IgG ICs; red channel: podocin. One-way ANOVA was used for statistical analyses in A and B. Lmer ANOVA was used for the statistical analyses in D-I. Significance levels are indicated with asterisks. *: p < 0.05, **: p < 0.01, ***: p < 0.001, ****: p < 0.0001.

**Figure 5.**
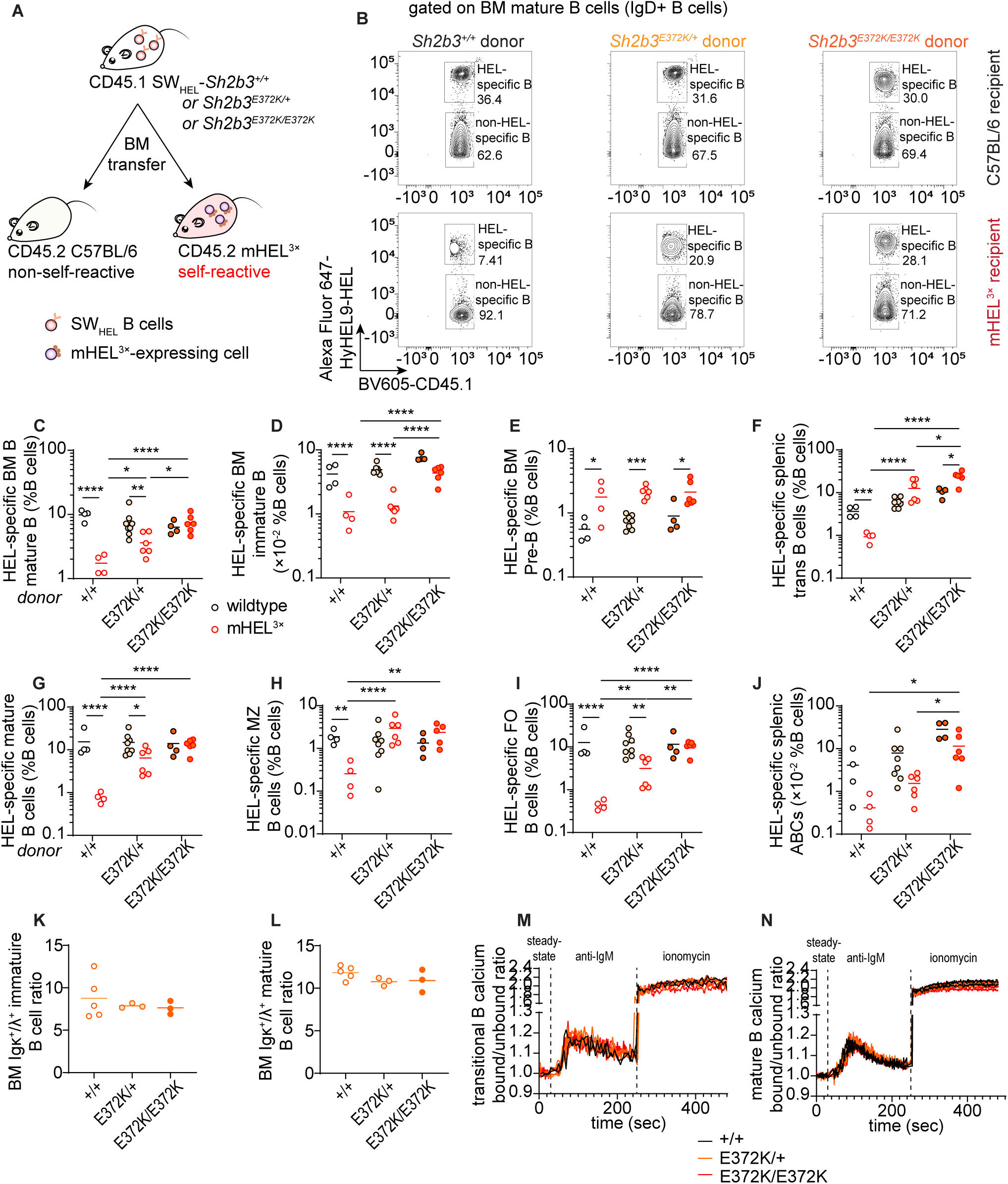
B cell tolerance breach in *Sh2b3^E372K^*mice. (**A**) Design of SW_HEL_- mHEL^3×^ BM chimera experiment for assessing B cell tolerance in *Sh2b3^+/+^*, *Sh2b3^E372K/+^*and *Sh2b3^E372K/E372K^* mice. (**B**) Representative flow cytometric plots showing the gating of HEL-specific (self-reactive) and non-HEL-specific (non-self-reactive) BM mature B cells in WT or mHEL^3×^ mice receiving BM from SW_HEL_-*Sh2b3^+/+^*, SW_HEL_-*Sh2b3^E372K/+^* or SW_HEL_-*Sh2b3^E372K/E372K^* donors. Dot plot demonstrating the frequencies of HEL-specific (**C**) BM mature B cells (**D**) BM immature B cells, (**E**) BM Pre-B cells, (**F**) splenic transitional B cells, (**G**) splenic mature B cells and (**H**) splenic MZ B cells, (**I**) splenic FO B cells and (**J**) splenic ABCs as percentages of total lymphocytes in WT or mHEL^3×^ recipient mice receiving BM from SW_HEL_-*Sh2b3^+/+^*, SW_HEL_-*Sh2b3^E372K/+^*or SW_HEL_- *Sh2b3^E372K/E372K^* mice. Ratios of BM Igκ^+^/Igλ^+^ (**K**) immature and (**L**) mature B cells in *Sh2b3^E372K^* mice. Calcium influx into the cytoplasm of peripheral blood (**M**) transitional and (**N**) mature B cells of *Sh2b3^E372K^* mice at steady state, following treatment with anti-IgM antibody and treatment with ionomycin measured by the ratios of bound/unbound indo-1 AM ester to Ca^2+^. Results are representative of two independent experiments. Means are shown as bars in C-L. Two-way ANOVA was performed on log_10_-transformed data in C-J due to meet the variance homoscedasticity. Significance levels are indicated with asterisks. One-way ANOVA was used for statistical analysis in C-L. *: p < 0.05, **: p < 0.01, ***: p < 0.001, ****: p < 0.0001.

**Figure 6.**
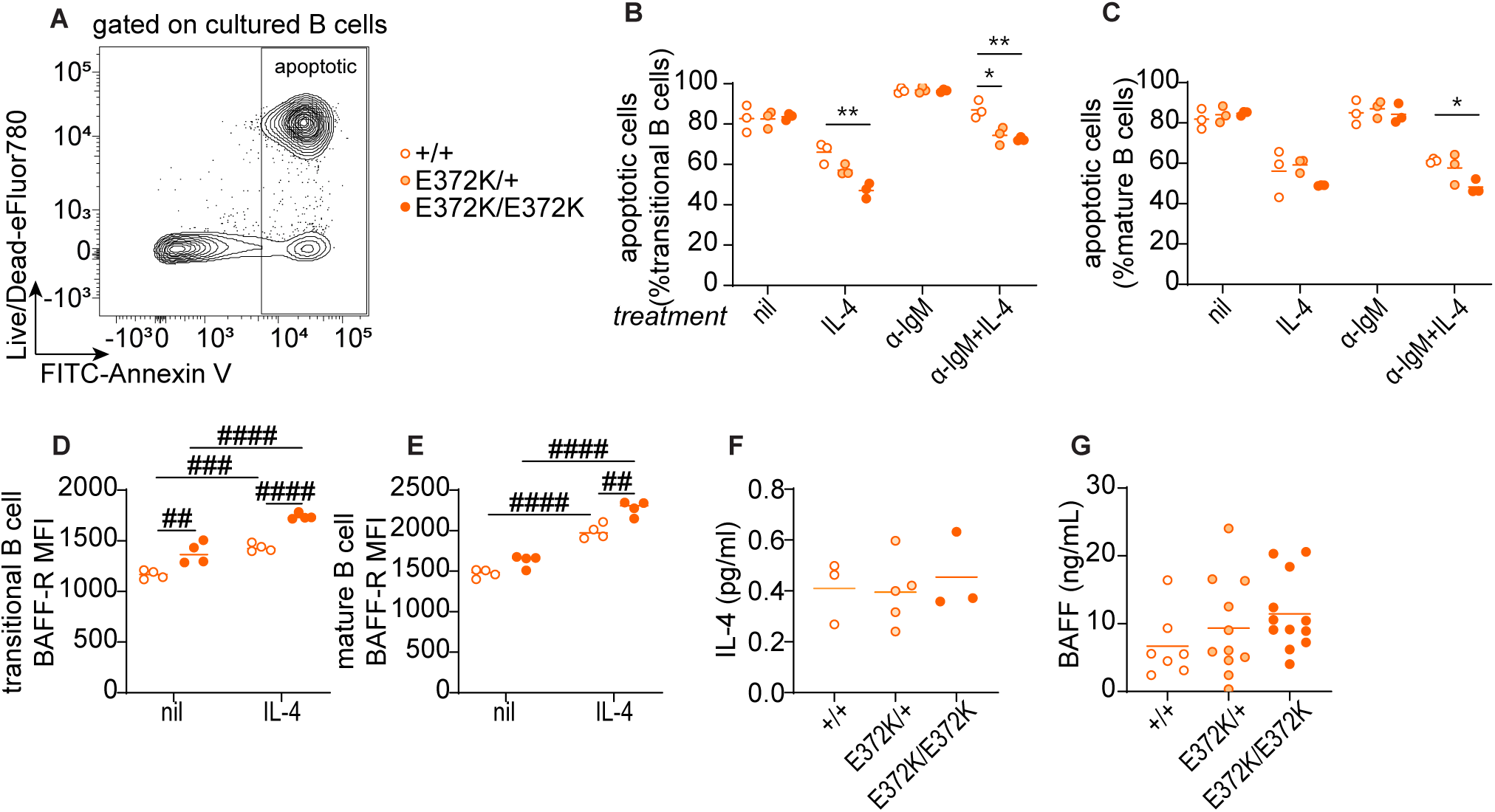
IL-4R and BAFF-R signaling in the survival in IgM-crosslinked splenic B cells of *Sh2b3^+/+^*, *Sh2b3^E372K/+^* and *Sh2b3^E372K/E372K^* mice. (**A**) Representative flow cytometry plot showing the gating strategy of apoptotic (Annexin^+^7-AAD^-/+^) cultured, sorted splenic B cells. Frequencies of apoptotic (**B**) transitional and (**C**) mature B cells in untreated, 20 ng/mL IL-4 only, 5 μg/mL anti-IgM (α-IgM) only, and 5 μg/mL α-IgM + 20 ng/mL IL-4 conditions. (**D**) BAFF-R expression measured by GMFI in (**D**) transitional and (**E**) mature B cells in untreated or 20 ng/mL IL-4 conditions. (**F**) Plasma IL-4 measured by Meso Scale Discovery assay. (**G**) Serum BAFF measured by bead-based immunoassay. Multiple unpaired student t tests were used for statistical analyses in B and C. Significance levels are indicated with asterisks. Two-way ANOVA was used for statistical analyses in D and E. Significance levels are indicated with hashes. One-way ANOVA was used for statistical analyses in F and G. */#: p < 0.05, **/##: p < 0.01, ***/###: p < 0.001, ****/####: p < 0.0001.

### *Sh2b3^E372K^* B cells are hyperresponsive to IL-4-induced rescue

IL-4 is known to exert anti-apoptotic effects on B cells during BCR-mediated negative selection (Granato et al., 2014). Since IL-4R signaling is in part transduced by JAK3 (Malabarba et al., 1995), which is regulated by SH2B3, we examined whether hypomorphic SH2B3 altered B cell apoptosis following BCR ligation. Treating transitional or mature B cells with IL-4 increased survival of B cells homozygous or heterozygous for the *E372K* allele to a greater extent than wild-type cells (**Figure A**-**C**). In contrast, IL-4 did not increase the survival of immature B cells isolated from BM (**Figure S 5S)**. Thus, augmented IL-4R signaling may contribute to defective negative selection in the spleen and thus the breach in B cell tolerance seen in *Sh2b3^E372K^* mice.

Another factor important for transitional B cell survival is BAFF-R signaling (Batten et al., 2000; Schiemann et al., 2001). We examined BAFF-R expression on splenic B cells following *in vitro* culture of transitional and mature B cells with and without IL-4, and were surprised to observe an increase in BAFFR expression (**Figure D, E, Figure S 5T and U**). This effect appeared to be greater in *Sh2b3^E372K/E372K^*and *Sh2b3^E372K/+^* cells, providing a possible mechanistic link between IL-4 signaling and accumulation of immature/transitional B cells in *Sh2b3^E372K/E372K^* and *Sh2b3^E372K/+^* mice. Serum concentrations of IL-4 and BAFF in *Sh2b3^E372K/E372K^*and *Sh2b3^E372K/+^* were similar to *Sh2b3^+/+^* (**Figure F and G**), suggesting that cellular phenotypes and the breach in tolerance are driven by receptor hyperresponsiveness or expression rather than an increase in available cytokines or growth factors.

## Discussion

In conjunction with previous work (Jiang et al., 2019), this study demonstrates that rare functional variants occurring in GWAS-associated SLE genes may contribute to disease pathogenesis. Often the strongest evidence for an association between a particular gene and a disease comes from GWAS – with many monogenic, disease-causing mutations occurring in genes that have also been identified by well-powered common variant associations. *SH2B3* has already been robustly linked to SLE by GWAS with the lead common variant (rs10774625) tagging a haplotype block that encompasses the whole of *SH2B3*, being a strong eQTL (Alcina et al., 2010; Bentham et al., 2015; Morris et al., 2016; Wang et al., 2021), and being in linkage with a relatively common missense *SH2B3* variant, rs3184504 (r2 = 0.92). Here, we add weight to the link between SH2B3 and SLE pathogenesis by showing that SLE patient-specific novel and ultrarare variants in *SH2B3* encode proteins with impaired function leading to dysregulated B cell development and tolerance that predisposes carriers to autoimmunity. Presence of the hypomorphic allele alone is insufficient to cause disease, consistent with occurrence in some unaffected family members. Instead, clinical disease is likely to require additional environmental or genetic triggers.

Five out of the six rare patient-specific variants impaired the ability of SH2B3 to suppress IFNγR signaling. Interestingly, the patient (D.II.1) who carried the A536T variant that retained normal suppressive function, was also compound heterozygous for *ACP5* (Supplementary table 2), which is a known cause of Spondyloenchondrodysplasia with immune dysregulation (SPENCDI) that shares features with SLE. In contrast, the *ADAR* variant found in patient A.II.1 is unlikely to be sufficient to cause SLE in the heterozygous state, and there may be an additional role for SH2B3.

Through structural studies, we showed that substitution of key residues responsible for forming the hydrogen-bonding network of the BC loop in the SH2 domain can result in reduced binding to target pTyr and domain instability. Indeed, our previous work on the SH2 domain of SH2B3 revealed similar examples of functional impairment, with V402M affecting more severely the binding to pTyr and inhibition of IFNγR signaling (Morris et al., 2021). Nevertheless, we were only able to co-crystallize murine SH2 domain of SH2B3 with pY813 JAK2 as expression of human SH2B3 was unsuccessful. In addition, we showed that patient-specific variants outside known functional domains (C133Y and R566Q) also impaired SH2B3 protein function. How these variants affect protein function will require future study on unannotated regions of the SH2B3 protein.

Using the SW_HEL_-mHEL^3×^ mouse model, we demonstrated incomplete negative selection of self-reactive *Sh2b3^E372K^*B cells at multiple immune checkpoints. This tolerance breach is also likely to be applicable in a non-transgenic setting since 55-75% of immature B cells have been shown to harbor autoreactivity in healthy individuals (Wardemann et al., 2003). In most individuals these autoreactive B cells are removed through tolerance checkpoint mechanisms, but this has been shown to be impaired in individuals with SLE (Yurasov et al., 2005). While evasion of normal tolerance mechanisms was not sufficient to cause spontaneous autoimmunity in *Sh2b3^E372/E372K^* mutant mice, sensitization with pristane treatment led to accelerated autoantibody production, suggesting that autoimmunity may occur in the presence of additional triggers.

Our data suggests that elevated IL-4R signaling and BAFF-R expression may be in part responsible for the impaired negative selection in the periphery. Engagement of BCR on developing B cells, can either induce negative selection or facilitate B cell maturation depending on the strength of BCR signal, as well as positive signals from other receptors including cytokine receptors, such as IL-4R and BAFF-R (Granato et al., 2014). Signaling through IL-4R provides pro-survival signals to BM immature and splenic transitional B cells by impeding the upregulation of Bim following BCR engagement (Granato et al., 2014). Consistently, we observed that IL-4 treatment of splenic transitional and mature wildtype B cells could rescue them from apoptosis following IgM-crosslinking, while the hyperresponsiveness of *Sh2b3^E372K/E372K^* B cells to IL-4 signaling appears to provide autoreactive B cells with higher survival potential. The binding of BAFF to BAFF-R supports the differentiation of T1 into T2 B cells and T2 B cell survival, as well as their further differentiation into mature naïve B cells and subsequent maintenance (Sasaki et al., 2004). Anergic autoreactive T1 B cells are more dependent on BAFF signaling due to elevated Bim expression, and thus compete less well for selection into the recirculating follicular pool, compared with non-autoreactive T1 B cells (Lesley et al., 2004). Therefore, enhanced IL-4R signaling and subsequent restrained Bim expression in autoreactive *Sh2b3^E372K/E372K^*B cells may mean they are not as dependent on BAFF as their wildtype counterparts. Unexpectedly, we also observed the elevation of surface BAFF-R expression on transitional B cells in *Sh2b3^E372K/E372K^* mice in cell culture, which may further increase their competitive survival advantage *in vivo*.

Interestingly, we have found that IL-4 treatment can upregulate BAFF-R expression in both splenic transitional and mature B cells. In contrast, IL-4 treatment has been reported to have no effect on BAFF-R expression on CD23^+^ immature B cells isolated from the BM (Granato et al., 2014), which may suggest that splenic transitional B cells respond differently to those in the BM. Regardless, our observations suggest that heightened IL-4R signaling in *Sh2b3^E372K/+^* and *Sh2b3^E372K/E372K^* mice may lead to increased BAFF-R expression and contributes to the breach of immune tolerance checkpoints. Notably BAFF was not higher in the serum of *Sh2b3^E372K/+^*or *Sh2b3^E372K/E372K^* mice, suggesting that any survival advantage is not a result of increased bioavailability of BAFF.

Increases in BAFF-R and IL-4R are unlikely to be the sole mechanisms that increase autoimmune susceptibility in carriers of *SH2B3* hypomorphic variants; we showed that IL-4 did not promote survival of immature B cells isolated from BM. Previous studies have suggested that overproduction of immature B cells in *Sh2b3^-/-^* mice is a consequence of dysregulated IL-7R and c-KIT signaling (Cheng et al., 2016; Takaki et al., 2000). Unrestrained IL-7R signaling promotes B cell precursor proliferation leading to increased B cell production in humans, (Parrish et al., 2009) and gain-of-function *IL-7R* mutations increase peripheral B cell numbers and contribute to precursor B-cell ALL (Almeida et al., 2021; Shochat et al., 2011). Other cytokine responses that may be enhanced in the absence of SH2B3 and also impair central and/or peripheral tolerance include IL-2 and IL-15, both of which promote the proliferation of activated B cells via IL-2R signaling (Armitage et al., 1995; Mingari et al., 1984). Additional candidates are IL-3, which supports the proliferation of murine (Palacios et al., 1984) and human B cell precursors (Crooks et al., 2000), as well as the survival and maturation of plasmacytoid dendritic cells; IL-6, which drives pristane-induced anti-DNA IgG and facilitates spontaneous GCs in autoimmune models (Richards et al., 1998); and IFN-γ, which induces T-bet expression in B cells and drives spontaneous GC formation in autoimmune models (Domeier et al., 2016; Jackson et al., 2016) and ABC formation (Naradikian et al., 2016; Peng et al., 2002).

Contrary to the leukopenia and lymphopenia observed in some SLE patients (Rivero et al., 1978), our SH2B3 mutant mouse strains exhibited increased lymphocytes. This is perhaps unsurprising since lymphopenia is usually related to disease activity in SLE (Lu et al., 2021; Sapartini et al., 2018) and mice with hypomorphic *Sh2b3* mutations do not spontaneously develop an autoimmune phenotype. Lymphocytosis is also observed in individuals with compromised lymphocyte apoptosis such as Autoimmune Lymphoproliferative Syndrome (ALPS) patients, and in the corresponding *Fas^lpr^* lupus-like mouse model (Adachi et al., 1996; Völkl et al., 2016). Our studies showed that hypomorphic *Sh2b3* can protect B cells from apoptosis *in vitro*, possibly preventing lymphopenia. Clinical investigations (Supplementary Table 1) indicated that only two of the seven SLE patients with *SH2B3* mutations in this study displayed lymphopenia.

Taken together, our study uncovers a role for SH2B3 in mouse and human B cell tolerance due to negative regulation of cytokine receptor signaling. It also provides a rationale for the use of JAK inhibitors (Jakinibs) already approved for several autoimmune conditions and currently in trials for SLE (Tanaka et al., 2022). The convergence between IL-4R signaling and BAFF-R expression in promoting B cell survival suggests that Jakinib therapy may not only restrain aberrant JAK-STAT signaling, but potentially dampen BAFF-R expression, limiting aberrant selection and/or survival of autoreactive B cells. Overall, this study highlights the importance of studying rare variants in SLE patients to understand mechanisms that drive or contribute to the loss of tolerance and disease pathogenesis, which may be targeted to prevent autoimmune development.

## Acknowledgments

We thank the personnel of the Australian Cancer Research Foundation Biomolecular Resource Facility (JCSMR) for Sanger sequencing; Jing Gao, Nikki Ross, Jenna Lowe, Nay Chi Khin and Lora Starrs for assistance with the generation of CRISPR mouse models; Robert Tunningley for assisting with an ADVIA experiment; staff of the Australian Phenomics Facility (APF) for care of experimental animals; Dr Harpreet Vohra, Michael Devoy and Cathy Gillespie from the Microscopy and Flow Cytometry Facility (MCRF), John Curtin School of Medical Research (JCSMR) for assistance with flow cytometry and imaging; Koula Diamand for technical assistance running the Mesoscale Discovery platform; Jessica Lovell of the Bruestle Group, JCSMR for assistance with cryosectioning; Dr Yafei Zhang of the APF NGS Team for WES of CRISPR mice; Pablo Fernández de Cañete Nieto for conjugation of HyHEL9-Alexa Fluor 647. Dr Teresa Neeman of the Biological Data Science Institute of the ANU provided consultation on the statistical analyses in this study. This research/project was undertaken with the assistance of resources and services from the National Computational Infrastructure (NCI) and National Collaborative Research Infrastructure Strategy (NCRIS) via Phenomics Australia (to J.E). This project is funded by Australian NHMRC Fellowship and program grants to GCV as well as funding from the Francis Crick Institute. This research project was undertaken with the assistance of computational resources and services from the National Computational Infrastructure (NCI) an NCRIS enabled capability supported by the Australian Government, and the Phenomics Translational Initiative funded by the Medical Research Future Fund.

## Author contributions

Supervision: J.I.E. and C.G.V.; Conceptualization: J.I.E., C.G.V., J.J.B., Y.Z. and R.M.; Investigation: Y.Z., R.M., A.M.D.L., X.M., N.J.K., P.K., G.J.B., J.Y.C., Q.S., H.W., C.M.T., T.L.H., M.S., Z.Y., F.B., A.C and D.A.F; Resources: G.B., V.A., J.C.L., A.H., P.T., D.M., J.T.F., G.D.W., M.S., M.J., M.C.C.; Writing: J.I.E., C.G.V. and Y.Z.; Funding acquisition: C.G.V.

## Materials and Methods

### Methods

#### Site-directed mutagenesis (SDM)

Primer pairs for site-directed mutagenesis were designed using Agilent’s primer design program (https://www.agilent.com/store/primerDesignProgram.jsp) following QuikChange XL II kit recommendations and ordered as 25 nmole DNA oligos (**Supplementary Table 4**) from Integrated DNA Technologies (Coralville, Iowa, USA). The oligos were resuspended at 100 μM in UltraPure DNase/RNase-free distilled water and then reconstituted to a working concentration of 125 ng/μL. DNA templates were reconstituted at 50 ng/μL. The reagents were mixed as per the QuikChange II XL Site-Directed Mutagenesis Kit instructions and loaded into a Mastercycler pro S (Eppendorf) with the settings recommended by the manual. After PCR, 1 μL of DpnI restriction endonuclease (New England Biolabs; cat # R0176L) was added to each reaction tube to digest DNA templates. The resulting products were transformed into DH5α competent E. coli cells for amplification, Sanger sequencing and selection.

#### Sanger sequencing

To prepare primers for Sanger sequencing, DNA oligos (**Supplementary Table 5**; Integrated DNA Technologies) were resuspended at 100 μM and then further diluted into 1.6 pmol/ μL (μM). Plasmid DNA templates were diluted to 250 ng/μL. The following reagents were mixed in each well of a 96-well PCR plate (Thermofisher Scientific; cat # AB-0600): 3.5 μL of 5× sequencing buffer, 1 μL BigDye Terminator 3.1 (Applied Biosystems; cat # 4337455), 1.6 pmol/μL sequencing primer, 5% (v/v) DMSO if the sequence was GC-rich and UltraPure DNase/RNase-free distilled water (Invitrogen; cat # 10977015) to a final volume of 20 μL. Thermocycling was performed with the following cycling conditions: 1 cycle of initial denaturation at 94°C for 5 min, 30 cycles of denaturation (96°C, 10 sec), annealing (50°C, 5 sec) and extension (60°C, 4 min), and hold at 4°C when all steps were complete. The resulting amplicons were purified using ethanol and EDTA before undergoing capillary sequencing at the Biomolecular Resource Facility (BRF), JCSMR. Results were analyzed using Sequencher (Gene Codes).

#### Cell culture and luciferase assay

HEK293 cells (ATCC; CRL-1573™) were cultured in complete DMEM (cDMEM) in Nunc EasYFlask T75 cell-culture flasks (Thermo Scientific; cat # 156499) and housed in CO_2_ incubator (5% CO_2_) at 37°C, with passaging every 3 days by a 1 in 10 split. All cell lines tested negative for mycoplasma contamination using PlasmoTest Mycoplasma Detection Kit (InvivoGen; cat # rep-pt1). To set up transfection for luciferase assays, HEK293 cells in 24-well plates were transfected with STAT1BS-Firefly (145 ng), pRL-CMV-Renilla (5 ng) and pReceiver-M12-*SH2B3* (100 ng) plasmids using Lipofectamine 2000 (Invitrogen; cat # 11668019) resuspended in Opti-MEM (Gibco; cat # 31985070) following the manufacturer’s instructions. The STAT1BS-GAS Firefly luciferase reporter is a gift from Vicki Athanasopoulos. Transfected cells were incubated for 24 hours before treatment with 50 ng/mL recombinant human IFN-γ (Peprotech; cat # 300-02) or cDMEM only control for 24 hours. Cell lysates were analyzed with Luc-Pair Duo-Luciferase assay kit (GeneCopoeia; cat # LF003) using a VICTOR Nivo multimode microplate reader (PerkinElmer; cat # HH35000500).

#### Expression and purification of LNK and SH2B SH2 domains

DNA encoding the mouse E372K and WT SH2B3 SH2 domain (residues 324– 446) and an N-terminal NusA fusion separated by a TEV cleavage in a pET-50b (+) was transformed into tuner (DE3) (Novagen; cat # 70623), BL21(DE3) *E. coli* cells and expression was induced by addition of 1 mM IPTG at 18 °C overnight. Cells were collected by centrifugation and frozen at −30 °C. Cells from 1 L culture were resuspended in 40 mL lysis buffer (20 mM Tris (pH 8.0), 10 mM imidazole (pH 8.0), 300 mM NaCl, 2 mM TCEP, 5 mM phenyl phosphate, 1 U DNase, 1 mM PMSF, and 20 mg lysozyme) containing EDTA-free protease inhibitor cocktail (Sigma-Aldrich; cat # 11697498001) and lysed by sonication. Lysate was clarified by spinning cells for 10 min at 20,000 × *g* before loading supernatant onto cOmplete™ His-Tag purification resin (Merck; cat # 5893682001). Bound proteins were washed with 20 mM tris (pH 8.0), 10 mM imidazole (pH 8.0) and 300 mM NaCl, 5 mM phenyl phosphate followed by 20 mM tris (pH 8.0), 30 mM imidazole (pH 8.0) and 300 mM NaCl, 2 mM TCEP 5 mM Phenyl Phosphate. Protein was eluted in 20 mM tris (pH 8.0), 250 mM imidazole (pH 8.0) and 300 mM NaCl, 5 mM Phenyl Phosphate and 2 mM TCEP. Eluate was then cleaved with TEV protease overnight at 4 °C and subsequently purified further by size exclusion chromatography (Superdex 200 26/600 from GE Healthcare; cat # GE28-9893-36) in TBS, 2 mM TCEP and 5 mM Phenyl Phosphate.

#### Thermostability assays

WT and E374K SH2B3 SH2 domains were desalted into 100 mM NaCl, 20 mM Tris (pH 8.0), 2 mM TCEP buffer and diluted to 100 μM. Where peptides were used, a five-fold molar excess of peptide was used, in phenyl phosphate conditions concentration was 8 mM. Each sample was transferred into a capillary and measured from 35 °C to 95 °C using a Tycho N6T (Nanotemper). Data were analyzed in Prism (GraphPad Software).

#### SPR assays

Direct binding experiments were performed on a Biacore 4000 (GE Healthcare) in 10 mM HEPES (pH 7.4), 150 mM NaCl, 3.4 mM EDTA, 0.005% Tween 20 using a streptavidin coated chip and were regenerated in 50 mM NaOH, 1 M NaCl. SH2B3 SH2 domains were flowed over a streptavidin coated chip for 420 s at 30 μL/min with immobilized biotinylated JAK2 pY813. A reference flow cell was included by passing buffer without protein over a single lane and the sensorgrams from the reference cell were subtracted from the experimental flow cell analyses. Data were subsequently plotted in Prism.

SPR competition assays were performed on a Biacore 4000 (GE Healthcare) in 10 mM HEPES (pH 7.4), 150 mM NaCl, 3.4 mM EDTA, 0.005% Tween 20 using a streptavidin coated chip and were regenerated in 50 mM NaOH, 1 M NaCl. A biotinylated peptide representing the IL6ST pY757 sequence was immobilized to the chip by passing over 1 μg/mL of peptide dissolved in 10 mM HEPES pH 7.4, 150 mM NaCl, 3.4 mM EDA, 0.005% Tween 20. 0.1–0.5 μM SH2B3 SH2 domain was pre-incubated phosphopeptides or phenylphosphate before being flowed over the chip for 240–720 seconds at 30 μL/min. Data was analyzed using Prism, the response of SH2B3 in the presence of the peptide was normalized to an SH2B3 only control and then fitted as an IC_50_ curve via non-linear regression.

#### Crystallography

The E372K SH2B3 SH2 domain was buffer exchanged into low salt buffer (20 mM Tris (pH 8.0), 2 mM TCEP and 100 mM NaCl) and crystal trays were set up with 5 mg/mL of protein and a 2-fold molar excess of peptide using vapor diffusion hanging drop experiments set up in house. The E372K SH2B3 SH2 domain crystallized in 18% PEG 8000, 0.05 M MgAc, 0.1 M Tris pH 8.5 and was cryo-protected in paratone and immediately snap frozen in liquid nitrogen. Data was collected at the MX2 beamline at the Australian Synchrotron. Data reduction, scaling and integration was performed using XDS. Crystal structure of the E372K SH2B3 SH2 domain was solved by molecular replacement (search model PDB ID: 7R8W) using Phaser as implemented in PHENIX(Liebschner et al., 2019). All structurers were refined using PHENIX and model building was performed in COOT(Emsley and Cowtan, 2004; Emsley et al., 2010). Structures were visualized using PyMOL(Schrodinger, 2015) (Schrödinger, Inc.).

#### Human patients and healthy controls

Written informed consent was obtained as part of the Australian Point Mutation in Systemic Lupus Erythematosus (APOSLE) study, the Centre for Personalised Immunology (CPI) program, and the Healthy Blood Donors (HBD) register (The Canberra Hospital, Garran, ACT, Australia). This study was approved by ANU and ACT Health Human Ethics Committees and was conducted in accordance with the guidelines of the National Statement on Ethical Conduct in Human Research (2007).

Whole blood of patients, relatives and healthy donors was collected by venipuncture at the antecubital area into 9 mL Vacutainer ACD-A collection tubes (BD; cat # 455055) and processed within 24 hr of collection. Peripheral blood mononuclear cells (PBMCs) were isolated by layering whole blood over Lymphoprep density gradient media (Stemcell Technologies; cat # 07851), resuspended in freezing media consisting of 90% fetal bovine serum (FBS) and 10% dimethyl sulfonate (DMSO), then stored in −80°C cryogenic freezer prior to analysis.

#### Mouse models

All mice were bred and housed in specific-pathogen-free (SPF) environments at the Australian Phenomics Facility (APF), Australian National University (ANU), Acton, ACT, Australia. All mouse-related procedures were approved by the Animal Experimentation Ethics Committees (AEEC) of the ANU. Mouse models carrying orthologous mutations of SLE patients were generated in C57BL/6 mice (Charles River Laboratories) via CRISPR-Cas9 genome editing technology according to Jiang *et al*. (Jiang et al., 2019) and Gurumurthy *et al*. (Gurumurthy et al., 2019). Three mouse models were generated:

*Sh2b3^E372K^*, *Sh2b3^Δ^* and *Sh2b3^R530Q^* (**Supplementary Table 6**). Briefly, single stranded oligonucleotides (ssODN) and single guide RNA (sgRNA) were purchased from IDT and delivered to the C57BL/6N^crl^ fertilized mouse zygotes as a ribonucleoprotein complex with the following concentrations: Cas9 protein (50 ng/µL), sgRNA (2.5 ng/µL) and ssODN (50 ng/µL). The edited mouse zygotes were surgically transferred into the uterine horn of the pseudopregnant CFW/crl mice. Founder mice were genotyped by Sanger sequencing using the primer pairs listed in the **Supplementary Table 6**. B6.129-*Rag1^tm1Mom^* (*Rag1^-/-^*) mice were used as BM recipients in our BM chimera experiments. C57BL/6-*Ptprc^a^*mice were used as wildtype congenic (CD45.1) BM donors in the BM chimera experiments. The mice were co-housed by litters and sexes with mixed genotypes.

#### 50:50 mixed BM chimera

BM cells were collected from CD45.1^+^ wildtype, CD45.2^+^-*Sh2b3^+/+^*and CD45.2^+^-*Sh2b3^E372K/E372K^* mice. Cells from CD45.1+ wildtype and CD45.2^+^-*Sh2b3^+/+^* or CD45.2_+_-*Sh2b3^E372K/E372K^* mice were mixed at 1:1 ratio, respectively, and 2 × 10^6^ BM cell mix injected into the tail vein of each corresponding *Rag1^tm1Mom^* (hereinafter referred to as Rag^-/-^) mouse that had received 500 cGy radiation from the RS2000 small animal irradiator (Rad Source Technologies). Injected mice were allowed 10 weeks for BM reconstitution before analysis.

#### SW_HEL_-mHEL^3×^ BM chimera

*Sh2b3^E372K/+^*(CD45.2) mice were crossed to SW_HEL_ mice (CD45.1) mice to generate *Sh2b3^+/+^*-SW_HEL_^Hc/-,Lc/-^ (CD45.1/2) and *Sh2b3^E372K/E372K^*-SW_HEL_^Hc/-,Lc/-^ (CD45.1/2) BM donor mice. Recipient membrane-bound HEL^3×^ (mHEL^3×^) mice (on CD45.1 congenic background) were a kind gift of Professor Robert Brink (Garvan Institute, Sydney, Australia). The mHEL^3×^ expressed in the mutant mice serve as a ubiquitously expressed self-antigen. BM was collected from donor mice and cryopreserved in freezing medium (90% FBS with 10% DMSO) at −80°C until needed. BM ablation of recipient mice was performed by double irradiation (450 cGy each time) 4 hr apart in a RS5000 small animal irradiator. Irradiated mice were injected with 2×10^6^ thawed donor BM cells and allowed for reconstitution for 8 weeks before analysis.

#### Pristane-induced murine SLE model

12-week-old *Sh2b3^E372K^* mice were injected with either 0.25 ml pristane or PBS intraperitoneally (randomly assigned). Retro-orbital blood was collected from injected mice before pristane injection and set periods thereafter to measure serum immunoglobulins and anti-nuclear antibodies (ANAs) by ELISA and assess cellular phenotypes. 6 months post-pristane injection mice were euthanized. Kidneys were either fixed on 10% formalin and stained for H&E to look at renal pathology or cryopreserved in OCT for immunofluorescence analysis.

#### Hematology analysis

200 μL mouse blood was collected via retro-orbital bleed into a Monovette 200 μL EDTA tube (Sarstedt; cat # 20.1288). Following collection, the blood samples were de-identified and assigned sample numbers during preparation and acquisition to achieve blinding. 100 μL of the anticoagulated blood was transferred into a Axygen 1.1 ml polypropylene cluster tubes (Corning; cat # MTS-11-C) containing 100 μL FACS wash. Samples were analyzed on an ADVIA 2120 hematology system (Siemens Healthineers).

#### IL-4-supplemented IgM crosslinking of B cells

Spleens were collected from *Sh2b3^+/+^* and *Sh2b3^E372K/E372K^*mice and meshed to generate single-cell suspension, which were stained with antibody cocktail containing BV480-CD93 (BioLegend; cat # 136507), PE-CD19 (BioLegend; cat # 115508), Alexa Fluor 647-B220 (BioLegend; cat # 103226) and Alexa Fluor 700-CD3 (BioLegend; cat # 100216) antibodies, then with 7-AAD (Invitrogen; cat # A1310). The cells were sorted into live (7-AAD^-^) transitional (CD19^+^CD3^-^ B220^+^CD93^+^) and mature B cells (CD19^+^CD3^-^B220^+^CD93^-^) on BD FACSAria Fusion and FACSAria II (BD), before cultured in complete RPMI-1640 media without/with 5 μg/mL goat anti-mouse IgM (Jackson ImmunoResearch; cat # 115-006-075) and/or recombinant murine 25 ng/ml IL-4 (Peprotech; cat # 214-14) for 16 hours at 37°C/5% CO_2_ before staining with fixable viability dye eFluor 780 (Invitrogen; cat # 65-0865-14) and FITC-Annexin V (BD; cat # 556419) for apoptosis analysis on a LSRFortessa X-20 flow cytometer (BD). To determine BAFF-R experssion, sorting and cell culture was performed in as described above for the apoptosis assay. Following 16-hr incubation at 37°C/5% CO_2_, the cultured B cells were stained with Fc block followed by fixable viability dye eFluor 780 and FITC BAFF-R monoclonal antibody (Invitrogen; cat #11-5943-81) and analyzed on a LSRFortessa X-20 flow cytometer (BD).

#### Flow cytometry

To prepare mouse spleens for flow cytometry analysis, collected spleens were isolated as single-cell suspensions. For BM cell isolation, claws were removed by incising the ankles of each leg, while muscles, ligaments and tendons were removed from the tibiae and femora with scalpel. Joints on one end of the cleaned tibiae and femora were cleaved. BM was collected with a 25G needle attached to a 1mL syringe. For preparing murine blood for flow cytometry antibody staining, plasma-separated mouse blood with anticoagulant (EDTA) was washed with FACS wash. To lyse red blood cells in these samples, RBC lysis buffer (168 mM NH_4_Cl, 10 mM KHCO_3_, 100 μM Na_2_EDTA, pH7.3) was added and incubated for 3-5 min before washing. The resulting cell suspension was enumerated and 3-5×10^6^ cells were plated in 96-well round bottom plates for staining. The samples were de-identified during collection and assigned numbers during preparation and acquisition to achieve blinding. Prior to antibody staining, cells were treated with TruStain FcX rat anti-mouse CD16/32 antibodies (BioLegend; cat # 101320) or human TruStain FcX CD16/32/64 antibodies (BioLegend; cat # 422302) to block Fc receptors. The cells were then stained with antibody cocktail (**Supplementary Table 7**) along with LIVE/DEAD Fixable Aqua Dead Cell Stain (Invitrogen; cat # L34957) or eBioscience Fixable eFluor780 viability dye (Invitrogen; cat # 65-0865-14). To fix and permeabilize samples, eBioscience Foxp3/transcription factor staining buffer set (Invitrogen; cat # 00-5523-00) was used per manufacturer’s instructions. Staining of intracellular markers were performed following permeabilization. Single-color controls and samples were acquired on a LSRFortessa X-20 flow cytometer (BD), while results were analyzed using FlowJo software (BD).

Flow cytometric analysis of human PBMCs was performed following methods described in the study by Ellyard *et al*. (Ellyard et al., 2019).

#### Calcium flux

Mouse peripheral blood was processed to remove RBC via lysis prior to Fc blocking and then stained with antibodies against surface markers including biotin-CD93 (Invitrogen, cat # 13-5892-85), BV605-CD19 (BD, cat # 115539), APC Cy7-CD3 (BioLegend, cat # 100222), Alexa Fluor 647-B220 (BioLegend, cat # 103226), and PE Cy7-streptavidin (BioLegend, cat # 405206). The cells were washed with complete RPMI-1640 pre-warmed to 37°C and stained with Indo-1 AM (Invitrogen, cat # I1223) diluted to 2 μM in warm RPMI-1640 and incubated at 37°C for 30 min before washed and stained with 7-AAD (Invitrogen, cat # A1310).

For acquisition, data scales of the detectors for the BUV395 (379/28) and BUV496 (515/30 450LP) filters were set to linear. Samples were kept at 37°C prior to and during acquisition. Baseline was recorded for 30 sec, followed by the addition of anti-mouse IgM (Jackson ImmunoResearch; cat # 115-006-075) to a final concentration of 10 μg/mL, immediately mixed before resuming acquisition for further 3.5 mins to measure the calcium flux following BCR ligation. At 4 mins post-acquisition, ionomycin (Invitrogen, cat # I24222) was added to the sample tube to a final concentration of 1 μg/mL, immediately mixed before before resuming acquisition for another 4 mins to measure maximal Ca^2+^ flux output.

#### Bead-based immunoassay

Serum BAFF quantitation was performed on plasma samples from 12-14 week-old *Sh2b3^E372K^* mice using the LEGENDplex Mouse B cell Panel – S/P (1-plex) w/VbP kit (BioLegend; cat # 740983) following the manufacturer’s instructions. Data were analyzed using the LEGENDplex Data Analysis Software Suite (BioLegend, https://www.biolegend.com/en-gb/legendplex/software).

#### Meso-Scale Discovery

Plasma samples from *Sh2b3^E372K^* mice were plated for the mouse U-plex Biomarker Group 1 Multiplex Assay Box 2 10-Assay Plate (Meso Scale Discovery, cat #K15322K) and the assay performed according to the manufacturer’s instructions. The plate was read on a Meso Sector S 600 (Meso Scale Discovery).

#### Treg Suppression Assay

Regulatory T cells (Tregs; CD4+CD25+), naïve T cells (CD4+CD62L+CD44lo) and antigen presenting cells (APC; CD3-) were purified by flow cytometry. Naïve T cells were labeled with Cell Trace Violet (ThermoFischer Scientific; cat # C34557) according to the manufacturer’s instructions. Naïve T cells (2×10^4^) were co-cultured with Tregs at the rations shown in the presence of APCs (4×10^4^ cells) and anti-CD3 (2μg/ml) for 72 hours at 37°C/5%CO_2_. T cell proliferation was measured by flow cytometry in the presence of 7AAD to allow for discrimination of live cells.

#### ELISA

Flat-bottom 96-well plates were pre-treated with 0.002% (w/v) poly-L-lysine solution for 5 hr at room temperature before coating with 50 μL of 50 ng/μL calf thymus DNA (Sigma-Aldrich; cat # D8661) or 5 ng/μL HEL, resuspended in ELISA coating buffer (50 mM Carbonate buffer, pH 9.6) overnight at 4°C in a humidified chamber. Coated plates were blocked with 150 μL of blocking buffer (1× PBS, 10% w/v BSA, 0.5v/v Tween20). Plasma samples diluted at 1:40 and 1:80 or positive controls (sera from MRL-*Fas* mice) diluted at 1:80, 1:160. 1:320, 1:640 and 1:1280 were added to the wells and incubated overnight at 4°C. The plates were washed and incubated with either AP-conjugated goat anti-mouse IgG or IgM antibody (SouthernBiotech; cat # 1030-04, 1020-04) (**Supplementary Table 8**), washed again and incubated with phosphatase substrate (Sigma-Aldrich; cat # S0942) dissolved in developing buffer (0.1 M glycine, 0.1 mM ZnCl_2_, 1.0 M MgCl_2_·6H_2_O). Absorbances at 405 nm and 605 nm (reference wavelength) were measured using an Infinite 200 Pro plate reader (Tecan).

The ANA ELISA was performed on sera collected from 17-week-old female *Sh2b3^E372K^* mice using a mouse anti-nuclear antigens (ANA/ENA), total immunoglobulins (IgA+IgG+IgM) ELISA kit (Alpha Diagnostic; cat # 5210) following manufacturer’s instruction.

#### Immunofluorescence (IF)

OCT-embedded, cryopreserved mouse kidneys were cut into 7-μm sections, fixed with acetone, blocked with blocking buffer (1× PBS, 3% BSA, 0.5% Triton X-100) and stained with goat anti-mouse Podocin antibody (Santa Cruz Biotechnology; cat # sc-22296) (**Supplementary Table 8**) and then with Alexa Fluor 594-conjugated donkey anti-goat IgG (H+L) and Alexa Fluor 488-conjugated donkey anti-mouse IgG (H+L) antibodies (Invitrogen; cat # A-21202). The stained sections were preserved in Vectashield antifade mounting medium with DAPI (Vector Laboratories; cat # H-1200-10) and imaged using an Olympus DP70 camera system attached to an IX70 fluorescence microscope (Olympus Life Science). A 200× magnification was used with 1/200 sec exposure for the green channel (IgG) and 1/50 sec exposure for the red channel (podocin). IgG IC deposition was scored blindly by the investigator following imaging.

#### Statistical analyses

One-way ANOVA using multiple comparison and Šidák correction was used for comparing differences among three or more groups of data from single experiments. Multiple student t-tests were used for comparing between two sets of data in each group of interest. Brown-Forsythe and Welch ANOVA tests were used for analyzing human immunophenotyping data. Student t-test was used for comparing the differences between CD45.2^+^/CD45.1^+^ cell percentage ratios in two types of mixed BM chimeras (wt:wt and hom:wt, of the *Sh2b3^E372K^* strain). Linear mixed-effects model (lmer) with estimated marginal means (emmeans) using experiment as a blocking factor was used for analyzing luciferase data. Linear mixed-effects model (lmer) ANOVA using experiment as a blocking factor was used for analyzing ADVIA and gross phenotype data pooled from multiple experiments. Fisher’s exact tests were used for analyzing categorical data in the IF experiment and glomerular scores. Variances of data from the SW_HEL_-mHEL^3×^ are not homoscedastic and therefore the data was log_10_-transformed before two-way ANOVAs to fulfill the assumption of homoscedasticity. All multiple student’s t-tests, one-way ANOVAs, two-way ANOVAs (for BM chimera data), Brown-Forsythe and Welch ANOVAs were performed in Prism (GraphPad Software). Two-way ANOVAs (for SW_HEL_-mHEL^3×^ data), lmer emmeans, lmer ANOVAs and Fisher’s exact tests were performed using R (R Foundation for Statistical Computing).

For testing the assumptions for each statistical test used, we followed the following scheme: in each one-way ANOVA, a Bartlett’s test of homogeneity of variance was used for testing the assumption of equal variances, a residual plot was generated to examine the fit for a linear model and a Q-Q plot was generated to assess the normality of data distribution; for two-way ANOVAs, residual plots and Q-Q plots were used for assessing the model assumptions; since Brown-Forsythe and Welch ANOVA tests does not assume equal variance but assumes normal distribution, Q-Q plots were generated to assess the normality of data distribution; lmer-based tests were utilized for analyzing pooled data from independent experiments to estimate and remove random effects due to variability introduced during independent experiment setups, residual plots were generated to assess the assumptions of constant variance and linearity of effects.

## Legends for Supplementary Figures

**Figure S1.**
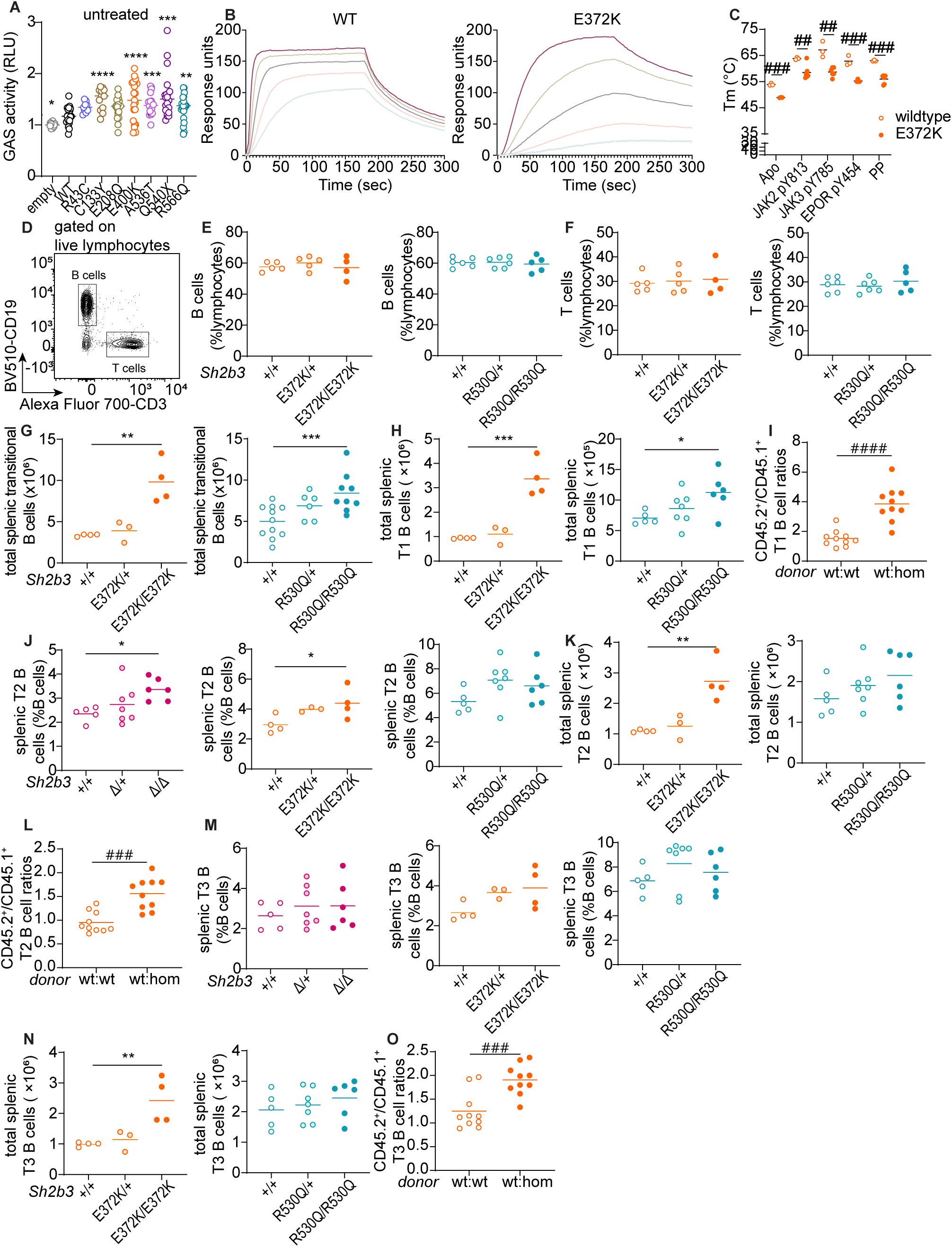
Human and murine SH2B3 protein functionality and gross phenotypes of *Sh2b3^Δ^*, *Sh2b3^E372K^* and *Sh2b3^R530Q^* mice and transitional B cell phenotypes in *Sh2b3^E372K^*and *Sh2b3^R530Q^* mice, and chimeras of *Sh2b3^E372K^* mice. (**A**) Relative GAS activity of unstimulated HEK-Blue IFN-α/β cells overexpressing wildtype or mutant human *SH2B3*. (**B**) Representative direct binding curves for the WT and E372K SH2B3 SH2 domains to a JAK2 pY813 peptide. (**C**) Melting temperatures for WT (unfilled) and E372K (filled) SH2B3 SH2 domains in their apo form and after the addition of the phosphomimetic phenyl phosphate (PP), JAK2 pY813, JAK3 pY785 and EPOR pY454 peptides. N = 3-7. Means are shown as bars. (**D**) Representative flow cytometric plot showing the gating strategy of B cells (CD19^+^CD3^-^) and T cells (CD19^-^CD3^+^). Dot plots showing frequencies of (**E**) B and (**F**) T cells as percentages of total splenic lymphocytes in *Sh2b3^E372K^* (upper panel) and *Sh2b3^R530Q^* (lower panel) mice. (**G**) Total numbers of splenic transitional B cells in *Sh2b3^E372K^*(left) and *Sh2b3^R530Q^* (right) mice. (**H**) Total numbers of splenic T1 B cells in *Sh2b3^E372K^* (left) and *Sh2b3^R530Q^* (right) mice. (**I**) CD45.2^+^/CD45.1^+^ T1 B cell ratios among splenic B cells in 50:50 BM chimeras of CD45.1-*Sh2b3^+/+^*and CD45.2-*Sh2b3^+/+^*/*Sh2b3^E372K/E372K^* mice. (**J**) Frequencies of T2 B cells as percentages of splenic B cells in *Sh2b3^Δ^* (left)*, Sh2b3^E372K^*(middle) and *Sh2b3^R530Q^* (right) mice. (**K**) Total numbers of splenic T2 B cells in *Sh2b3^E372K^* (left) and *Sh2b3^R530Q^*(right) mice. (**L**) CD45.2^+^/CD45.1^+^ T2 B cell ratios among splenic B cells in 50:50 BM chimeras of CD45.1-*Sh2b3^+/+^*and CD45.2-*Sh2b3^+/+^*/*Sh2b3^E372K/E372K^* mice. (**M**) Frequencies of T3 B cells as percentages of splenic B cells in *Sh2b3^Δ^*(left), *Sh2b3^E372K^* (middle) and *Sh2b3^R530Q^* (right) mice. N) Total numbers of splenic T3 B cells in *Sh2b3^E372K^* and *Sh2b3^R530Q^* mice. Frequencies of (**O**) CD45.2^+^/CD45.1^+^ T3 B cell ratios among splenic B cells in 50:50 BM chimeras of CD45.1-*Sh2b3^+/+^*and CD45.2-*Sh2b3^+/+^*/*Sh2b3^E372K/E372K^*mice. Results in A are pooled from seven independent experiments. Data in E, F and G are pooled from 2-4 independent experiments. Results in H-O are representative of 2-3 independent experiments. Means are indicated as bars. Lmer ANOVAs using experiment as a blocking factor were used for statistical analyses in A (all conditions were compared to cells transfected with WT SH2B3), D and E. Student-t tests were used for the statistical analysis in C, I, L and O. One-way ANOVA was used for the statistical analyses in E-H, J, K, M and N. Significance levels in lmer ANOVAs are indicated with asterisks, while those in multiple student-t tests are indicated with hashes. */#: p < 0.05, **/##: p < 0.01, ***/###: p < 0.001, ****/####: p < 0.0001.

**Figure S2.**
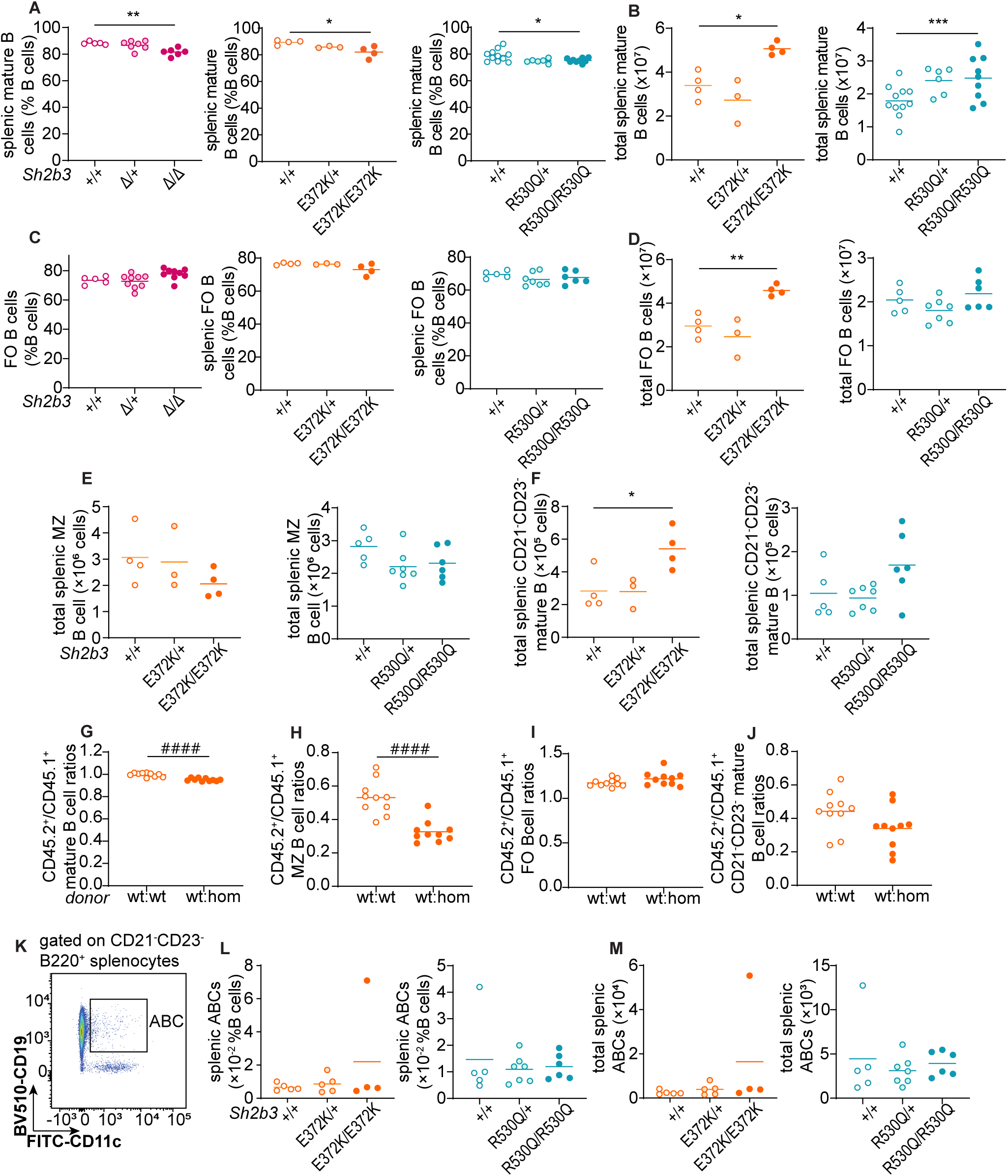
Mature B cell phenotypes in *Sh2b3^Δ^*, *Sh2b3^E372K^* and *Sh2b3^R530Q^* mice, and chimeras of *Sh2b3^E372K^*mice. (**A**) Frequencies of mature B cells as percentages of splenic B cells. (**B**) Total numbers of splenic mature B cells. (**C**) Frequencies of FO B cells as percentages of splenic B cells. (**D**) Total splenic FO B cells. Total splenic (**E**) MZ B cells and (**F**) CD21^-^CD23^-^ mature B cells. CD45.2^+^/CD45.1^+^ ratios of frequencies of splenic (**G**) mature B cells, (**H**) MZ B cells, (**I**) FO B cells and (**J**) CD21/35^-^CD23^-^ mature B cells in 50:50 BM chimeras of CD45.1-*Sh2b3^+/+^* and CD45.2-*Sh2b3^+/+^*/*Sh2b3^E372K/E372K^*mice. **K**. Representative flow cytometric plot of splenic atypical memory B cells (ABCs; B220^+^CD21/35^-^CD23^-^CD11c^+^CD19^+^). (**L**) Frequencies and (**M**) total numbers of splenic ABCs. Results are representative of 2-3 independent experiments. Means in A-F, L and M are shown as bars. One-way ANOVA was used for statistical analysis of immunophenotyping data (A-F, L and M) while student t-test was used for analyzing data from BM chimera experiments (G-J). Significance levels of one-way ANOVAs are indicated with asterisks while those of student-t tests are indicated with hashes. Significance level criteria are indicated as follow: */#: p < 0.05, **/##: p < 0.01, ***/###: p < 0.001, ****/####: p < 0.0001.

**Figure S3.**
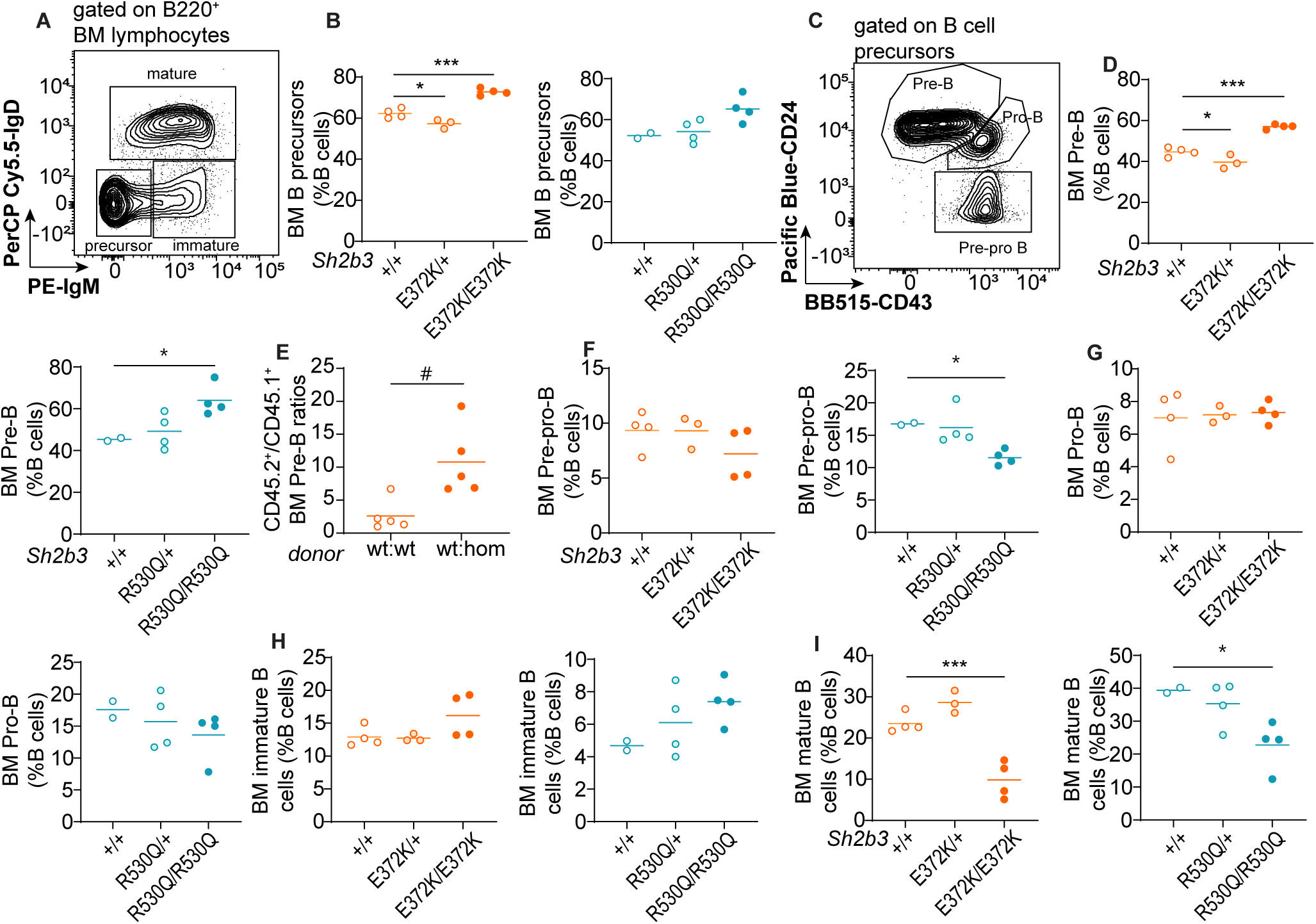
BM B cell phenotypes in *Sh2b3^E372K^* and *Sh2b3^R530Q^* mice. **A-I**. BM B cell phenotypes. (**A**) Representative flow cytometric plot showing the gating of B cell precursors (IgM^-^IgD^-^), immature (IgM^+^IgD^-^) and mature (IgD^+^) B cells. Cells were pregated on B220^+^ lymphocytes. (**B**) Frequencies of B cell precursors as percentages of BM B cells. (**C**) Representative flow cytometric plot showing the gating of pre-pro (CD24^-^CD43^+^), pro-(CD24^hi^CD43^+^) and pre-B cells (CD24^+^CD43^-/lo^). Frequencies of pre-B cells as percentages of BM B cells in (**D**) *Sh2b3^E372K^* (left) and *Sh2b3^R530Q^* (right) mice and (**E**) 50:50 BM chimeras of of CD45.1-*Sh2b3^+/+^* and CD45.2-*Sh2b3^+/+^*/*Sh2b3^E372K/E372K^*mice. Frequencies of (**F**) pre-pro and (**G**) pro-, (**H**) immature and (**I**) mature cells as percentages of BM B cells. Lines in dot plots show means and results are representative of two independent experiments. One-way ANOVA was used for statistical analysis of immunophenotyping data (B, D, F-M) while student t-test was used for analyzing data from BM chimera experiments (E). Significance levels of one-way ANOVAs are indicated with asterisks while those of student t-tests are indicated with hashes. Significance level criteria are indicated as follow: */#: p < 0.05, **/##: p < 0.01, ***/###: p < 0.001, ****/####: p < 0.0001.

**Figure S4.**
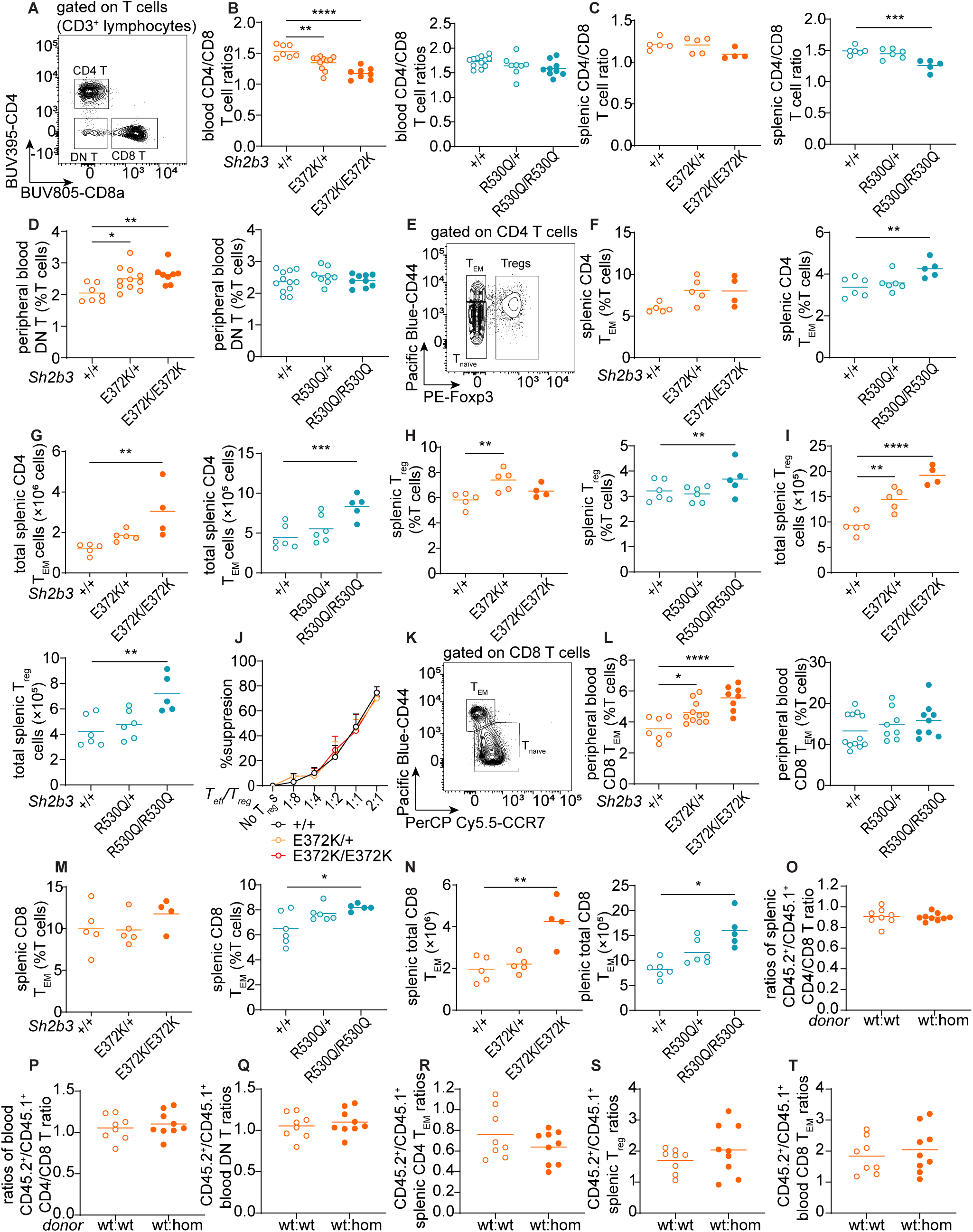
Peripheral blood and splenic T cell phenotypes in *Sh2b3^E372K^* and *Sh2b3^R530Q^*mice. (**A**) Representative flow cytometric plot showing the gating of double-negative (DN; CD4^-^CD8^-^), CD4^+^ (CD4^+^CD8^-^) and CD8^+^ (CD4^-^ CD8^+^) T cells. Dot plots showing the ratios of (**B**) blood and (**C**) splenic CD4/CD8 T cells in *Sh2b3^E372K^* mice and splenic CD4/CD8 T cells in *Sh2b3^R530Q^* mice. (**D**) Frequencies of DN T cells in the peripheral blood of *Sh2b3^E372K^* mice. (**E**) Representative flow cytometric plot showing the gating of CD4 naïve T (T_naïve_: CD44^lo/-^Foxp3^-^), effector memory T (T_EM_: CD44^hi^Foxp3^-^) and regulatory T (T_reg_: Foxp3^+^) cells. (**F**) Frequencies and (**G**) total numbers of CD4 T_EM_ cells in the spleens of *Sh2b3^E372K^*and *Sh2b3^R530Q^* mice. (**H**) Frequencies and (**I**) total numbers of T_reg_ cells in the spleen of *Sh2b3^E372K^* (left) and *Sh2b3^R530Q^* (right) mice. (**J**) Percentage suppression of effector T cells (T_eff_) by T_reg_s in culture at various T_eff_/T_reg_ ratios by using T_reg_s sorted from the spleens of *Sh2b3^+/+^*, *Sh2b3^E372K/+^* and *Sh2b3^E372K/E372K^* mice. (**K**) Representative flow cytometric plot showing the gating of CD8 T_naïve_ (CD44^lo/-^ CCR7^lo^) and T_EM_ (CD44^hi^CCR7^-^) cells. Frequencies of (**L**) peripheral blood and (**M**) splenic CD8 T_EM_ cells in *Sh2b3^E372K^* and *Sh2b3^R530Q^* mice. (**N**) Total numbers of splenic CD8 T_EM_ in *Sh2b3^E372K^* and *Sh2b3^R530Q^*mice. Ratios of CD45.2^+^/CD45.1^-^ (**O**) splenic and (**P**) peripheral blood CD4/CD8 T ratios, (**Q**) peripheral blood DN T cells, (**R**) splenic T_EM_ cells, (**S**) T_reg_s and (**T**) peripheral blood CD8 T_EM_ cells in 50:50 BM chimeras of CD45.1-*Sh2b3^+/+^*and CD45.2-*Sh2b3^+/+^*/*Sh2b3^E372K/E372K^*mice. (**T**) Results are representative of two independent experiments in B-D, F-I and L-N and those in J and O-T are from a single experiment. One-way ANOVA was used for statistical analysis of immunophenotyping data (B-D, F-I and L-N) while student-t test was used for analyzing data from BM chimera experiments (O-T). Significance levels of one-way ANOVAs are indicated with asterisks while those of student-t tests are indicated with hashes. Significance level criteria are indicated as follow: */#: p < 0.05, **/##: p < 0.01, ***/###: p < 0.001, ****/####: p < 0.0001.

**Figure S5.**
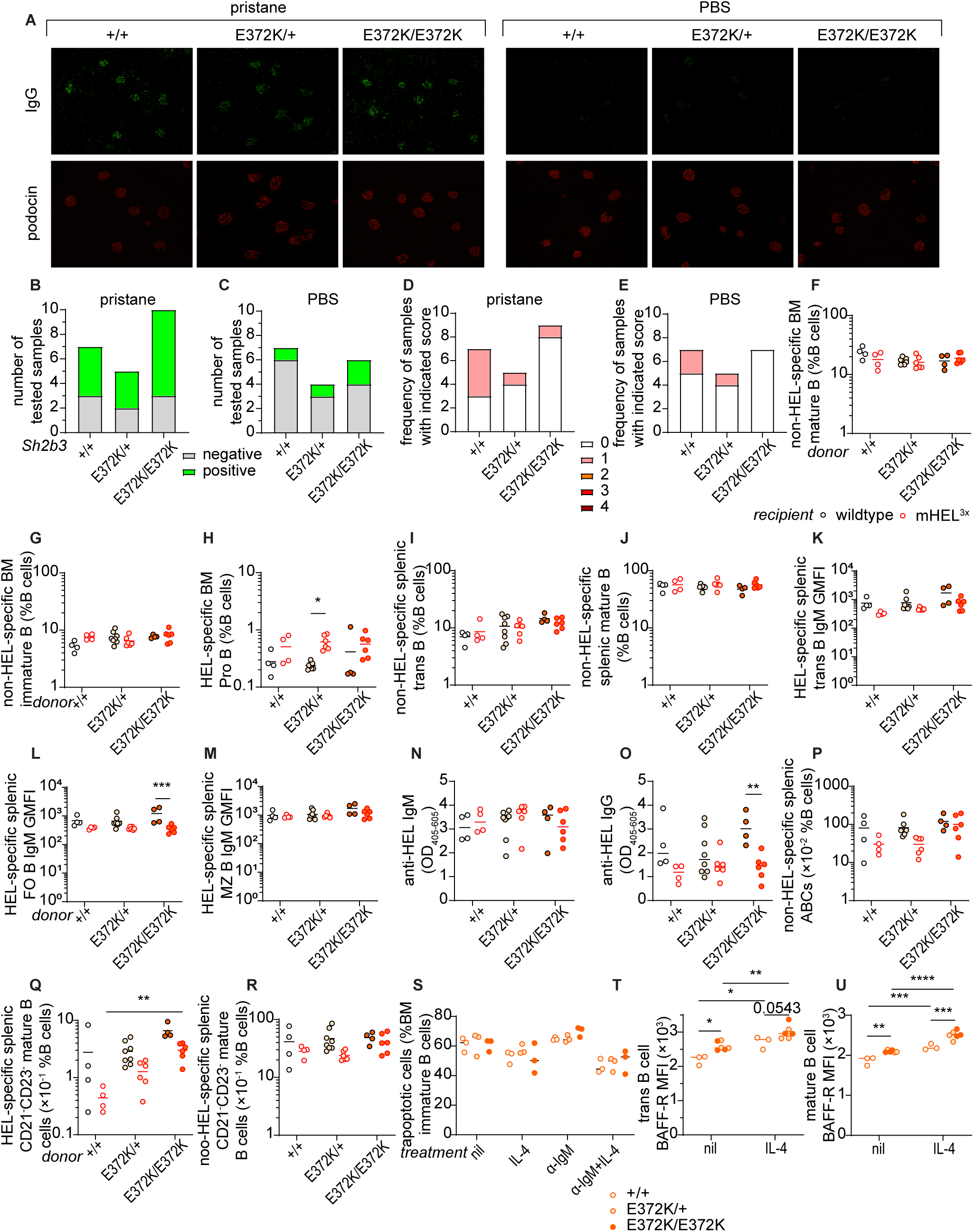
Spontaneous autoimmunity and B cell tolerance in *Sh2b3^E372K^* mice. **A-C**. IgG immune-complex deposits in mice treated with PBS or pristane. (**A**) Representative immunofluorescence images showing IgG (green) and podocin (red) staining in the kidney sections of pristane/PBS-treated *Sh2b3^E372K^* mice 20 weeks following treatment with (**B**) pristane and (**C**) PBS and numbers of kidney sections that are positive and negative for IgG ICs in the glomeruli of *Sh2b3^E372K^* mice treated with (**B**) pristane and (**C**) PBS. **D-E.** Numbers of kidney sections with indicated glomerular score 20 weeks following treatment with (**D**) pristane and (**E**) PBS. **G-R**. Cellular and serological phenotypes of SW_HEL_-mHEL^3×^ chimeric mice. Frequencies of non-HEL-specific BM (**F**) mature and (**G**) immature B cells, (**H**) HEL-specific Pro-B cells, non-HEL-specific splenic (**H**) transitional and (**I**) mature B cells, relative surface IgM expression on HEL-specific splenic (**J**) transitional, (**L**) FO and (**M**) MZ B cells in WT (black circles) or mHEL^3×^ (red circles) recipients receiving BM from SW_HEL_-*Sh2b3^+/+^* (unfilled), SW_HEL_-*Sh2b3^E372K/+^* (yellow filling) or SW_HEL_-*Sh2b3^E372K/E372K^* (orange filling) donors. Levels of anti-HEL (**N**) IgM and (**O**) IgG in wildtype and mHEL^3×^ mice receiving BM from SW_HEL_-*Sh2b3^+/+^*, -*Sh2b3^E372K/+^* and -*Sh2b3^E372K/E372K^* donors measured as OD_405-605_ by ELISA. Frequencies of (**P**) non-HEL-specific ABCs, (**Q**) HEL-specific and (**R**) non-HEL-specific CD21^-^ CD23^-^ mature B cells as percentages of splenic lymphocytes. (**S**) Percentages of apoptotic BM immature B cells in untreated, 20 ng/mL IL-4 only, 5 μg/mL anti-IgM (α-IgM) only, and 5 μg/mL α-IgM + 20 ng/mL IL-4 conditions. BAFF-R surface expression on (**T**) transitional and (**U**) mature B cells measured by flow cytometry as median fluorescence intensity (MFI). Data in A-C is pooled from two independent experiments. Results in D-U are representative of 2-3 independent experiments. Fisher’s exact test was used for the statistical analyses in B-E. Means in D-U are shown as bars, two-way ANOVA was used for statistical analysis in G-U. Significance levels for two-way ANOVAs are indicated with asterisks. *: p < 0.05, **: p < 0.01, ***: p < 0.001, ****: p < 0.0001.

## Supplementary Table Legends

**Supplementary Table 1**. Diagnosis and clinical manifestations of probands carrying SH2B3 variants.

**Supplementary Table 2**. Rare variants in genes known to cause human SLE in the probands identified through whole exome sequencing.

**Supplementary Table 3**. Data collection and refinement statistics for the SH2 domains of murine SH2B3 protein with phosphopeptides bound.

**Supplementary Table 4**. List of primers used for introducing patient-specific variants and other published variants into mammalian expression vectors of human *SH2B3*.

**Supplementary Table 5**. Sequencing primers designed for validating ORF sequences in the mammalian expression vectors of human *SH2B3* via Sanger sequencing.

**Supplementary Table 6**. List of guide RNAs (gRNAs), single-stranded oligodeoxynucleotides (ssODNs) for CRISPR/Cas9 gene editing of mouse models, and oligos for validating the editing results by Sanger sequencing.

**Supplementary Table 7**. Antibodies, fluorochrome-conjugated streptavidin and viability dyes used in flow cytometry.

**Supplementary Table 8**. Primary and secondary antibodies used for ELISA and immunofluorescence.

## Abbreviations

ABC: **a**typical memory **B c**ell
ALPS: **A**utoimmune **L**ympho**p**roliferative **S**yndrome
ANA: **a**nti-**n**uclear **a**ntibody
BAFF-R: **B** cell **a**ctivating **f**actor **r**eceptor
BM: **b**one **m**arrow
FO B: **fo**llicular **B** (cell)
HEL: **h**en **e**gg **l**ysozyme
IC: **i**mmune **c**omplex
IFNγR: **I**nter**f**ero**n gamma r**eceptor
IL-4R: **I**nter**l**eukin **4 r**eceptor
mHEL^3×^: **m**embrane-bound **HEL** mutant carrying **three** substitutions
MZ B: **m**arginal **z**one **B** (cell)
PB: **p**lastma**b**last
pTyr: **p**hospho-**tyr**osine
SH2: **S**rc **h**omology **2** (domain)
SH2B3: **SH2B** adaptor protein **3**
SLE: **s**ystemic **l**upus **e**rythematosus
SWHEL: **sw**itch **HEL**
T1/2/3 (B cells): **t**ransitional **1**/**2**/**3** (B cells)
IC: **i**mmune **c**omplex
SH2B3: **SH2B** adaptor protein **3**
SLE: **s**ystemic **l**upus **e**rythematosus

