## Supplementary Tables 1-8 for "Rare *SH2B3* coding variants identified in lupus patients impair B cell tolerance and predispose to autoimmunity"

**Supplementary Table 1: Sex, ethnicity, onset, diagnosis and clinical manifestations of probands carrying *SH2B3* variants**

| Patient | Variant | Sex | Ethnicity | Age of SLE onset | Diagnosis | Clinical history |
|---|---|---|---|---|---|---|
| **A.II.1** | C133Y | F | Caucasian/ European | 15 | SLE with lupus nephritis, secondary immunodeficiency (hypogammaglobulinemia) and coagulopathy | **Postnatal**: dysmorphic features, failure to thrive, slow feeding habits, and physical delays.<br>**At the age of 15**: presented with fever, numerous spontaneous hematomas on the extremities and petechiae, swelling of the knees, polyarthralgias, frequent diarrheal stools accompanied by abdominal pain, inappetence and weight loss. Elevated erythrocyte sedimentation rate, pancytopenia, lymphopenia, coagulation disorder (prolonged prothrombin and activated partial thromboplastin time), proteinuria (0.64 g in 24-hr urine sample). Positive ANA in high titer (1:51200), positive anti-dsDNA, anti-histone and antiphospholipid antibodies (aPL), anti-cardiolipin (aCL) and lupus anticoagulant (LAC)) and low C3, C4 and CH50. Other laboratory findings included hypovitaminosis D3 and B12 and a high titer of antibodies to tissue transglutaminase. Renal biopsy was not performed. Enlarged liver with signs of diffuse lesion and splenomegaly was noticed on abdominal ultrasound, hypertrophic synovium of the elbows and knees with increased Doppler flow (synovitis) on diagnostic ultrasound of the locomotor system and retinal detachment of the left eye with maculopathy (dystrophy) of the right eye. She soon developed grand mal type epileptic seizures, without MRI signs typical for vasculitis of the central nervous system. The patient was treated with glucocorticoids (she received six intravenous pulses of glucocorticoids, followed by oral glucocorticoids) with hydroxychloroquine and azathioprine. She received antiepileptic drugs to establish satisfactory seizure control. |

Continued on next page

Supplementary Table 1 – continued from previous page

| Patient | Variant | Sex | Ethnicity | Age of SLE onset | Diagnosis | Clinical history |
|---|---|---|---|---|---|---|
| | | | | | | Because of hypogammaglobulinemia, she required replacement therapy with IVIG and because of severe CD4+ lymphopenia prophylaxis for opportunistic infections. Later she developed osteoporosis with multiple vertebral fractures (treated with bisphosphonates), arterial hypertension and recurrent episodes of gastroenterocolitis accompanied by dehydration and metabolic acidosis. |
| **B.I.2** | E208Q | F | Caucasian European | 33 | SLE, DVT | History of Hashimoto's thyroiditis.<br>**At the age of 33:** diagnosis of SLE after longstanding history of mild photosensitive rashes and mild intermittent small joint arthralgias and sicca syndrome which was managed with hydroxychloroquine 200-400 mg daily alone. The patient was tested positive for ANA with a homogeneous serum ANA pattern. History of pulmonary emboli, deep vein thrombus and 2$\times$ miscarriage. Other manifestations include alopecia, ulcerative pharyngitis, headache, dry eyes and aptyalism.<br>**At the age of 48:** diagnosed with antiphospholipid syndrome when borderline anticardiolipin antibody was detected. APLS was managed with regular warfarin.<br>**At the age of 62:** Patient self-ceased plaquenil due to concern re visual floaters. |

Continued on next page

Supplementary Table 1 – continued from previous page

| Patient | Variant | Sex | Ethnicity | Age of SLE onset | Diagnosis | Clinical history |
|---|---|---|---|---|---|---|
| **C.II.1** | E400K | M | Caucasian English | 77 | SLE, DVT | **At the age of 73:** diagnosis of rheumatoid arthritis with small joint arthralgias, trialled on methotrexate.<br>**At the age of 77:** diagnosis of SLE, developed noninfectious, nonmalignant pleural effusion and noted ANA >2560 and elevated dsDNA antibody. Scl70 positive on ENA testing. Other manifestations include presence of anticardiolipin antibody, low serum C4, ulcerative pharyngitis, Raynaud's disease, lymphocytopenia and photosensitivity.<br>**At the age of 79:** diagnosed with bilateral DVT, treated with warfarin. SLE managed with a combination of low dose methotrexate and prednisone, complicated by multiple skin cancers and cellulitis. Hydroxychloroquine commenced 2010 (at the age of 87), methotrexate ceased late 2010. |
| **D.II.2** | A536T | M | Chinese/ Asian | 9 | SLE, ITP, KD | **At the age of 6**:Diagnosed with ITP (presented with epistaxis and bruises), treated with IVIG with good resolution of symptoms.<br>**At the age of 6**: presented with fever, rash, cervical lymphadenopathy, conjunctivitis and mucosal involvement and diagnosed with Kawasaki Disease. Treated with 2 courses of IVIG. Developed coronary artery involvement as part of Kawasaki Disease, experienced intermittent episodes of second-degree heart block (Mobitz type 1 with Wenckebach phenomenon).<br>**At the age of 9**: SLE diagnosed – treated with Methotrexate<br>**At the age of 10**: Presented with nephrotic syndrome and blood in urine and decreased kidney function – kidney biopsy showed class IV lupus nephritis – treated with steroids and mycophenolate with good response.<br>**At the age of 12**: Developed low mood, poor concentration and diagnosed with cerebral lupus – treated with IV methylprednisolone with resolution of the symptoms. |

Continued on next page

Supplementary Table 1 – continued from previous page

| Patient | Variant | Sex | Ethnicity | Age of SLE onset | Diagnosis | Clinical history |
|---|---|---|---|---|---|---|
| **E.II.1** | Q540X | M | Hispanic, English/ European | 16 | SLE, ITP, DVT | **At the age of 16**: presented with severe SLE with ITP and secondary APS with DVT, Class IV glomerulonephritis, as well as skin and joint involvement. Positive anti-dsDNA, anti-Sm and anti-$\beta$2GPI antibodies, low serum complement. The patient was treated with cyclophosphamide, and maintained on 2000 mg/day mycophenolate, 400 mg/day hydroxychloroquine and warfarin. |
| **E.II.3** | Q540X | F | Hispanic, English/ European | - | $ANA^{+}$ | **At the age of 26**: Moderate titre of antinuclear antibodies (1:320), speckled pattern. Antibodies to extractable nuclear antigens, CCP, phospholipids and double-stranded DNA not detected. Circulating immunoglobulins and complements were normal. No proteinuria. Patient also reported symptoms consistent with Raynaud's phenomenon affecting the hands and the feet, and patches of alopecia. |
| **F.II.1** | R566Q | F | English | 29 | SLE with lupus nephritis | **At the age of 29**: presented with rash and polyarthritis, associated with renal involvement, elevated anti-dsDNA (1600 IU/ml), and positive anti-Ro60 antibodies. Also diagnosed with coexistent autoimmune thyroid disease with hypothyroidism, positive TPO antibodies. Renal biopsy revealed diffuse proliferative glomerulonephritis, for which she received 6 cycles of iv cyclophosphamide, maintained on a combination of mycophenolate mofetil, plaquenil and low dose prednisolone. She maintained clinical remission from later in the year onwards, but remained subclinically active with low C3, C4 and low-level proteinuria for many years; all of which have gradually improved. |

Continued on next page

Supplementary Table 1 – continued from previous page

| Patient | Variant | Sex | Ethnicity | Age of SLE onset | Diagnosis | Clinical history |
|---|---|---|---|---|---|---|
| **G.II.2** | R566Q | F | European | 12 | SLE | **At the age of 12**: presented with acute lupus featuring joint and muscle problems, required blood transfusions for a haemolytic anaemia. Rental biopsy revealed glomerulonephritis. The patient was initially treated with mycophenolate, then with intravenous cyclophosphamide pulses for six months, which put her into remission and the treatment reverted to mycophenolate. The patient had high dose prednisolone until the age of 17. Despite remission, the patient had ongoing metacarpophalangeal joint and foot pain and swelling, as well as intermittent mouse ulcers.<br>**At the age of 17**: normal joints, no evidence of skin rashes, serologically in remission (normal complements, anti-dsDNA and ANAs), although experiences recurrent sinusitis and tonsillitis. The patient is still on mycophenolate. |

**Supplementary Table 2: Rare variants in genes known to cause human SLE in the probands.**

| Gene | A.II.1 | B.I.2 | C.II.1 | D.II.1 | E.II.1 | F.II.1 | G.II.2 |
|---|---|---|---|---|---|---|---|
| *C1QA* | | | | | | | |
| *C1QB* | | | | | | | |
| *C1QC* | | | | | | | |
| *C1R* | | | | | | | |
| *C1S* | | | | | | | |
| *C2, CFB* | | | | | | | |
| *C3* | | | | | | | |
| *C4A* | | | | | | | |
| *C4B* | | | | | | | |
| *DNASE1* | | | | | | | |
| *TREX1* | | | | | | | |
| *PRKCD* | | | | | | | |
| *DNASE1L3* | | | | | | | |
| *ACP5* | | | | p.G109R[1]<br>p.C238R[2] | | | |
| *TNFSF6* | | | | | | | |
| *IFIH1* | | | | p.A542E | | | |
| *SOCS1* | | | | | | | |
| *NCKAP1L* | | | | | | | |
| *SAMHD1* | | | | | | | |
| *ADAR1* | p.I939V | | | | | | |
| *RNASEH2B* | | | | | | | |
| *TMEM173* | | | | | | | |

[1] ClinVar pathogenic variant
[2] Novel variant

**Supplementary Table 3: Data collection and refinement statistics for the SH2 domains of murine SH2B3 protein with phosphopeptides bound**

| E372K SH2B3 SH2 + JAK2 pY813 | |
|---|---|
| **Data Collection** | |
| Space group | C 2 |
| Cell dimensions<br>a, b, c (Å)<br>$\alpha$, $\beta$, $\gamma$ (°) | <br>75.82 37.61 50.49<br>90 108.76 90 |
| $R_{meas}$ (%) | 4.93 (76.49) |
| I /sigma(I) | 20.45 (2.49) |
| $CC_{1/2}$ (%) | 99.90 (81.80) |
| Completeness (%) | 99.04 (97.32) |
| Redundancy | 6.8 (6.9) |
| **Refinement** | |
| Resolution (Å) | 34.53 - 1.64<br>(1.70 - 1.64) |
| No. reflections | 111266 (11000) |
| R-work | 17.9 (23.8) |
| R-free | 21.4 (26.9) |
| No. atoms | |
| protein | 914 |
| solvent | 76 |
| B factors | |
| protein | 32.45 |
| solvent | 43.76 |
| r.m.s. deviations | |
| RMS(bonds) | 0.006 |
| RMS(angles) | 0.85 |
| Ramachandran favored (%) | 95.41 |
| Ramachandran allowed (%) | 4.59 |
| Ramachandran outliers (%) | 0.00 |
| Rotamer outliers (%) | 2.08 |

**Supplementary Table 4: List of primers used for introducing patient-specific variants and other published variants into mammalian expression vectors of human *SH2B3*.** Base change in each primer is in **red**.

| Oligo Name | Sequence (5' → 3') |
|---|---|
| h*SH2B3*sdmR43CFW | CGGGAGCTGGCC**T**GCCAGTACTGGC |
| h*SH2B3*sdmR43CRV | GCCAGTACTGGC**A**GGCCAGCTCCCG |
| h*SH2B3*sdmC133YFW | GCCCGGGCCCT**A**CTCCTTCCAGC |
| h*SH2B3*sdmC133YRV | GCTGGAAGGAG**T**AGGGCCCGGGC |
| h*SH2B3*sdmE208QFW | GCCTGGCCGAC**C**AGGCCTCCATG |
| h*SH2B3*sdmE208QRV | CATGGAGGCCT**G**GTCGGCCAGGC |
| h*SH2B3*sdmE400KFW | GACGCGGCGTGGG**A**AATACGTGCTCACT |
| h*SH2B3*sdmE400KRV | AGTGAGCACGTATT**T**CCCACGCCGCGTC |
| h*SH2B3*sdmA536TFW | CGCCCGAAGAACTG**A**CCAACAGCCTGCAG |
| h*SH2B3*sdmA536TRV | CTGCAGGCTGTTGG**T**CAGTTCTTCGGGCG |
| h*SH2B3*sdmQ540XFW | GGCCAACAGCCTG**T**AGCACCTGGAGCA |
| h*SH2B3*sdmQ540XRV | TGCTCCAGGTGCT**A**CAGGCTGTTGGCC |
| h*SH2B3*sdmR566QFW | CCGGAGCCACCTGC**A**GGCCATAGACAATC |
| h*SH2B3*sdmR566QRV | GATTGTCTATGGCC**T**GCAGGTGGCTCCGG |

**Supplementary Table 5: Sequencing primers designed for validating ORF sequences in the mammalian expression vectors of human *SH2B3* via Sanger sequencing.**

| Oligo Name | Sequence (5' → 3') |
|---|---|
| h*SH2B3*seqF1 | CTCCTCGCCCTCTTCC |
| h*SH2B3*seqF2 | GAAGTTCCTGCCCTGG |
| h*SH2B3*seqF3 | CATTTCCTGTCCTGCTACC |
| h*SH2B3*seqF4 | CAGAGGGTCTCCCAGG |
| h*SH2B3*seqR | CTGCAGCGACACCAG |

**Supplementary Table 6: List of guide RNAs (gRNAs), single-stranded oligodeoxynucleotides (ssODNs) for CRISPR/Cas9 gene editing of mouse models, and oligos for validating the editing results by Sanger sequencing**

| Oligo Name | Sequence (5' $\rightarrow$ 3') |
|---|---|
| $Sh2b3^{R530Q}$ gRNA | ACTGGTTGTCAATGGCCCGA |
| $Sh2b3^{R530Q}$ ssODN | AAATGGTTCCATTCACACGTCTGCCTCTCTGCACAGCTGTGAGAG AGGGGTGTACTGGTTGTCAATGGCC**T**GA**AGG**TGGCCCCGTGAAGA GGAGTCCATGTCATAGTCCGAGTCCCGGGCACTGCTCACAGACTC GAGCTC |
| $Sh2b3^{E372K}$ gRNA | TGAGTACATACTCTCCTCTC |
| $Sh2b3^{E372K}$ ssODN | CCCCACCCTGGTCCACCAGTTCCTGCCCCTACCTTGGCTCTGCCC TGTAAGTTGAATGTGAGTACATACT**T**TCCTCTC**CGG**GACTCACTC TGCCGCACCAGGAACACGCCGTGGGCATCAGGGCCCTGGAGCTGG ACCAGC |
| $Sh2b3^{\Delta}$ gRNA | TGAGTACATACTCTCCTCTC |
| m$Sh2b3^{R530Q}$seqFW | CTTGGTCTCTGGGTGTCCTT |
| m$Sh2b3^{R530Q}$seqRV | GCACTGTCCACGCTCTGT |
| m$Sh2b3^{E372K}$seqFW | CTTGGTCTCTGGGTGTCCTT |
| m$Sh2b3^{E372K}$seqRV | GCACTGTCCACGCTCTGT |

**Supplementary Table 7: Antibodies, fluorochrome-conjugated streptavidin and viability dyes used in flow cytometry.**

| Reactivity | Antigen /dye | Fluorochrome/ conjugate | Clone | Manufacturer | Cat. no |
|---|---|---|---|---|---|
| mouse | CD16/32 (FcX) | purified | 93 | BioLegend | 101320 |
| anti-HEL Ab | HEL (Ag) | purified | - | - | - |
| mouse | CD93 | biotin | AA4.1 | Invitrogen (eBiosciences) | 13-5892-85 |
| mouse | CD138 | biotin | 281-2 | BD Pharmingen | 553713 |
| mouse | CXCR5 | biotin | 2G8 | BD Pharmingen | 551960 |
| mouse | B220 | biotin | RA3-6B2 | BioLegend | 103204 |
| mouse | B220 | BUV395 | RA3-6B2 | BD Horizon | 563793 |
| mouse | CD4 | BUV395 | GK1.5 | BD Horizon | 563790 |
| mouse | CD19 | BUV395 | 1D3 | BD Horizon | 563557 |
| mouse | CD23 | BV421 | B3B4 | BioLegend | 101621 |
| mouse | I-A/I-E (MHCII) | BV421 | M5/114.15.2 | BioLegend | 107631 |
| mouse | PD-1 | BV421 | 29F-1A12 | BioLegend | 135218 |
| phosphatidylserine | Annexin V | Pacific Blue | - | BioLegend | 640918 |
| mouse | CD24 | Pacific Blue | M1/69 | BioLegend | 101820 |
| mouse | CD44 | Pacific Blue | IM7 | BioLegend | 103020 |
| amine | Live/dead fixable aqua | - | - | Invitrogen | L34965 |
| mouse | CD93 | BV480 | AA4.1 | BD Optibuild | 746239 |
| mouse | CD19 | BV510 | eBio1D3 | Invitrogen (eBiosciences) | 56-0193-82 |
| mouse | CD95 | BV510 | Jo2 | BD Horizon | 563646 |
| biotin | Streptavidin | BV510 | - | BioLegend | 405234 |
| phosphatidylserine | Annexin V | FITC | - | BD Pharmingen | 556419 |

Continued on next page

Supplementary Table 7 – continued from previous page

| Reactivity | Antigen /dye | Fluorochrome/ conjugate | Clone | Manufacturer | Cat. no |
|---|---|---|---|---|---|
| mouse | BAFF-R | FITC | eBio71-122-E16 | Invitrogen (eBio-science) | 11-5943-81 |
| mouse | CD11b | FITC | M1/70 | BD Pharmingen | 7222717 |
| mouse | CD11c | FITC | N418 | BioLegend | 117306 |
| mouse | CD62L | FITC | H1.2F3 | BD Pharmingen | 553236 |
| mouse | Foxp3 | FITC | FJK-16s | Invitrogen (eBio-sciences) | 2007700 |
| mouse | IgM | FITC | II/41 | BD Pharmingen | 553437 |
| mouse | CD34 | BB515 | MEC14.7 | BioLegend | 119302 |
| mouse | BST2 (CD317) | PE | 927 | Biolegend | 127009 |
| mouse | CD3 | PE | 17A2 | BioLegend | 100206 |
| mouse | CD4 | PE | RM4-5 | eBioscience | 12-0042-83 |
| mouse | CD25 | PE | PC61 | BioLegend | 102008 |
| HEL | HyHEL9 | PE | HyHEL9 | prepared by P.F. Cañete | - |
| mouse | Foxp3 | PE | FJK-16s | Invitrogen (eBio-sciences) | 4323635 |
| mouse | IgD | PE | 11-26c-2a | BioLegend | 405706 |
| mouse | Ig$\lambda$ | PE | RML-42 | BioLegend | 407308 |
| mouse | PD-1 | PE | J43 | Invitrogen (eBio-science) | 4293188 |
| mouse | CD21/35 | BV605 | 7G6 | BD Horizon | 583176 |
| mouse | CD43 | BV605 | S7 | BD Horizon | 563205 |
| mouse | CD45.1 | BV605 | A20 | BioLegend | 110737 |
| mouse | CD86 | BV605 | GL1 | BD Horizon | 563055 |
| mouse | CD138 | BV605 | 281-2 | BioLegend | 142515 |
| biotin | Streptavidin | BV605 | - | BioLegend | 405229 |
| mouse | CD23 | BV711 | B3B4 | BD Horizon | 563987 |
| DNA | 7-AAD | 7-AAD | - | Invitrogen | A1310 |
| mouse | CCR7 | PerCP-Cy5.5 | 4B12 | BioLegend | 120116 |

Continued on next page

| Reactivity | Antigen /dye | Fluorochrome/ conjugate | Clone | Manufacturer | Cat. no |
|---|---|---|---|---|---|
| mouse | CD4 | PerCP-Cy5.5 | RM4-5 | BioLegend | 100540 |
| mouse | IgD | PerCP-Cy5.5 | 11-26c.2a | BD Pharmingen | 564273 |
| mouse | CD4 | PE-Cy7 | RM4-5 | BD Pharmingen | 552775 |
| mouse | CD98 | PE-Cy7 | RI.388 | BioLegend | 128214 |
| mouse | IgD | PE-Cy7 | 11-26c.2a | eBioscience | 25-5993-82 |
| mouse | IgM | PE-Cy7 | II/41 | Invitrogen (eBioscience) | 25-5790-81 |
| mouse | KLRG1 | PE-Cy7 | 2F1 | eBioscience | 25-5893-82 |
| biotin | Streptavidin | PE-Cy7 | - | BioLegend | 405206 |
| mouse | CD45.1 | APC | 104 | BioLegend | 109814 |
| mouse | SiglecH | APC | 551 | BioLegend | 129611 |
| mouse | Bcl6 | Alexa Fluor 647 | K112-91 | BD Pharmingen | 561525 |
| mouse | CD4 | Alexa Fluor 647 | RM4-5 | BioLegend | 100530 |
| mouse | CD11c | Alexa Fluor 647 | N418 | BioLegend | 117312 |
| mouse | CD25 | Alexa Fluor 647 | PC61 | BioLegend | 102020 |
| HEL | HyHEL9 | Alexa Fluor 647 | HyHEL9 | Prepared by P. F. Cañete | - |
| amine | Fixable viability dye eFluor780 | eFluor780 | - | Invitrogen (eBioscience) | 65-0865-14 |
| mouse | CD3 | Alexa Fluor 700 | 17A2 | BioLegend | 100216 |
| mouse | CD4 | Alexa Fluor 700 | RM4-5 | BD Pharmingen | 557956 |
| mouse | CD19 | Alexa Fluor 700 | eBio1D3 | Invitrogen (eBioscience) | 56-0193-82 |
| mouse | Ig$\kappa$ | Alexa Fluor 700 | RMK-45 | BioLegend | 409508 |

Continued on next page

Supplementary Table 7 – continued from previous page

| Reactivity | Antigen /dye | Fluorochrome/ conjugate | Clone | Manufacturer | Cat. no |
|---|---|---|---|---|---|
| mouse | CD45.2 | BUV737 | 104 | BD Horizon | 564880 |
| mouse | CD8a | BUV805 | 53-6.7 | BD Horizon | 564920 |
| human | FcX | purified | 3G8 (CD16), FUN-2 (CD32), 10.1 (CD64) | BioLegend | 422302 |
| human | IgM | eFluor450 | SADA4 | Invitrogen (eBiosciences) | 48-9998 |
| human | IgD | BV510 | IA6-2 | BioLegend | 348220 |
| human | CD24 | BV605 | MLS | BioLegend | 311124 |
| human | CD19 | BV605 | HIB19 | BioLegend | 302238 |
| human | CD38 | PerCP-Cy5.5 | HIT2 | BD Pharmingen | 551400 |
| human | IgA | PE | IS11-8E10 | Miltenyi Biotech | 130-093-128 |
| human | CD10 | PE-CF594 | HI10a | BD Horizon | 562396 |
| human | IgG | PE-Cy7 | G18-145 | BD Pharmingen | 561298 |
| human | CD21 | APC | B-ly4 | BD Pharmingen | 561767 |
| human | CD27 | APC-eFluor780 | O323 | Invitrogen (eBiosciences) | 47-0279 |

**Supplementary Table 8: Primary and secondary antibodies used for ELISA and immunofluorescence.**

| Host | Reactivity | Antigen | Conjugate | Clone | Manufacturer | Cat. no |
|---|---|---|---|---|---|---|
| **ELISA** | | | | | | |
| goat | mouse | IgM | human ads-AP | polyclonal | Southern Biotech | 1020-04 |
| goat | mouse | IgG | human ads-AP | polyclonal | Southern Biotech | 1030-04 |
| **Immunofluorescence** | | | | | | |

Continued on next page

Supplementary Table 8 – continued from previous page

| **Host** | **Reactivity** | **Antigen** | **Conjugate** | **Clone** | **Manufacturer** | **Cat. no** |
|---|---|---|---|---|---|---|
| goat | mouse | podocin | purified | C-18 | Santa Cruz Biotechnology | sc-22296 |
| donkey | mouse | IgG (H+L) | Alexa Fluor 488 | polyclonal | Invitrogen | A-21202 |
| donkey | goat | IgG (H+L) | Alexa Fluor 594 | polyclonal | Invitrogen | A-11058 |
